# Epigenetic mechanisms governing cell type specific somatic expansion and toxicity in Huntington’s disease

**DOI:** 10.1101/2025.05.21.653721

**Authors:** Matthew Baffuto, Kert Mätlik, Isaac Ilyashov, Hasnahana Chetia, Maria Esterlita Siantoputri, Erika Sipos, Yurie Maeda, Paul Darnell, Nicholas Didkovsky, Laura Kus, Thomas S. Carroll, Douglas Barrows, Christina Pressl, Nathaniel Heintz

**Affiliations:** Laboratory of Molecular Biology, The Rockefeller University, New York, NY, United States of America; Bioinformatics Resource Center, The Rockefeller University, New York, NY, United States of America

## Abstract

Huntington’s disease (HD) is characterized by neuronal dysfunction and degeneration that varies markedly by brain region and cell type. Using high-resolution epigenetic profiling of postmortem human cell types we identify a pathogenic cascade linking cell type specific enhancer activity to somatic CAG expansion, and toxicity to epigenetic dysregulation. Enhancers regulating mismatch-repair (MMR) gene expression explain the specificity of expansion. In the second, toxic phase of HD we identify two distinct epigenetic mechanisms that disrupt regulation of hundreds of genes in the majority of HD MSNs, including several that cause haploinsufficient neurological disorders. Together, these data unify enhancer function, impaired DNA demethylation, and transcriptional dysregulation into a single model highlighting therapeutic opportunities that combine inhibition of somatic CAG expansion with restoration of neuronal DNA demethylation.

## Main text

Huntington’s disease (HD) is a late onset, autosomal dominant, neurodegenerative disease which results in motor and cognitive deficits, eventually resulting in death ^1^. In the initial stages of HD, somatic expansion of the CAG repeat in exon 1 of the mutant Huntingtin gene (*mHTT*) occurs in specific structures ^2^ and cell types in the human brain ^3–5^. Studies in humans and mouse models have demonstrated that genes involved in DNA mismatch repair (MMR) modulate the HD age of onset ^6–8^ and the capacity for somatic CAG expansion of the expanded HTT alleles (emHTT) in the mouse central nervous system (CNS) ^9–12^. In the human striatum, somatic CAG expansion in medium spiny neurons (MSNs) coincides with high levels of nuclear DNA mismatch repair complex, MutSb (MSH2/MSH3) ^5^. Although these data suggest that the high levels of MutSb in MSNs drive somatic instability in the striatum, the mechanisms regulating its abundance in MSNs or governing the cell type specific expression and function of other key MMR genes in HD have not been determined. As HD progresses, cell types carrying unstable *mHTT* alleles with somatically expanded CAG repeats have severe HD-associated transcript level changes ^3–5^. These findings support a model in which a second step of HD pathogenesis is caused by disruption of transcriptional regulation leading to cellular dysfunction and loss ^13^. Studies in HD mouse models and human tissue have demonstrated that transcriptional dysregulation is accompanied by a variety of epigenetic changes, including mild epigenetic aging ^14^, downregulation of cell identity genes ^15,16^, changes in post-translational histone modifications (PTMs) ^17,18^, reversal of polycomb-dependent developmental repression ^19^, and disease locus specific alterations of higher order chromatin structure ^20^. Which of these mechanisms occur specifically in human neurons in HD is not known.

Here we have used high resolution epigenetic analysis of cell types in the striatum, cerebral cortex, hippocampus, and cerebellum to understand the specificity of somatic expansion in the HD brain and to identify mechanisms of toxicity in cell types carrying somatically expanded *mHTT* CAG repeats. The finding that MutSb is elevated in human MSNs, coupled with the demonstration that there are cell type-specific enhancers in *MSH2*, *MSH3*, *FAN1* and other genes involved in somatic CAG expansion, suggests that selective somatic expansion of the *mHTT* CAG repeat in human cell types is driven by cell type specific enhancer engagement controlling the expression of these critical DNA repair genes. We demonstrate also that chromatin accessibility changes in HD brain are cell type specific, and that their magnitude and number is maximal in cell types in which the *mHTT* CAG repeat is unstable. We identify three distinct epigenetically regulated classes of genes in MSNs that are distinguished by differential enhancer and gene body accessibility, H3K27ac abundance and DNA methylation/hydroxymethylation in CpG (mCG/hmCG) or non-CpG (mCH/hmCH) dinucleotides. The magnitude, scale, and cell-specificity of epigenetic changes in neurons undergoing somatic CAG expansion, and the observation that genes responsible for haploinsufficient neurological or neurodegenerative diseases (i.e *PDE8B, ATP2B1, HIVEP2* etc) are strongly repressed in striatal MSNs suggest that the mechanisms driving cell type and neuronal circuit specific toxicity in HD arises from dysregulation of essential cellular functions in the majority of MSNs. These data both support a new mechanistic model of HD pathogenesis and identify new targets for HD therapy.

### Cell type specific transcriptional control of somatic expansion in the human brain

Based on the elevated expression of *MSH2* and *MSH3*, the abundance of their encoded proteins in MSN nuclei, and their pro-expansion functions in mismatch DNA repair, we have proposed that the relative abundance of MMR proteins in the nucleus may determine their propensity for somatic CAG expansion ^5,8–10^. However, the mechanisms responsible for this elevated expression have not been identified. Given that accessibility and enhancer function have been well documented for their contribution to cell-type-specific gene expression, we mapped candidate cis-regulatory elements (cCREs) using ATAC-seq profiles from FANS sorted nuclear populations ^4,5,21^ in: neurons showing extensive *mHTT* CAG expansion in Huntington’s disease (striatonigral (dMSN) and striatopallidal (iMSN) projection neurons, motor cortex derived L5a, L6a, and L6b pyramidal cells); neurons with moderate or minimal expansion, (hippocampal CA1 and CA2/3 neurons, ADARB2+ interneurons, dentate granule cells and cerebellar granule cells); glia which display negligible expansion (striatal oligodendrocytes, microglia, and astrocytes) (Fig 1A, fig S4b). Consistent with previous murine and human studies ^22,23^, UMAP analysis of consensus ATAC-seq peak sets generated by an iterative peak-calling strategy ^24^ separated neurons from glia and further distinguished individual neuronal and glial subtypes (Fig 1B). Correlation matrix clustering confirmed these relationships, demonstrating strong within-cell-type correlations and similarities among related neuronal populations (fig S1b).

**Fig 1.**
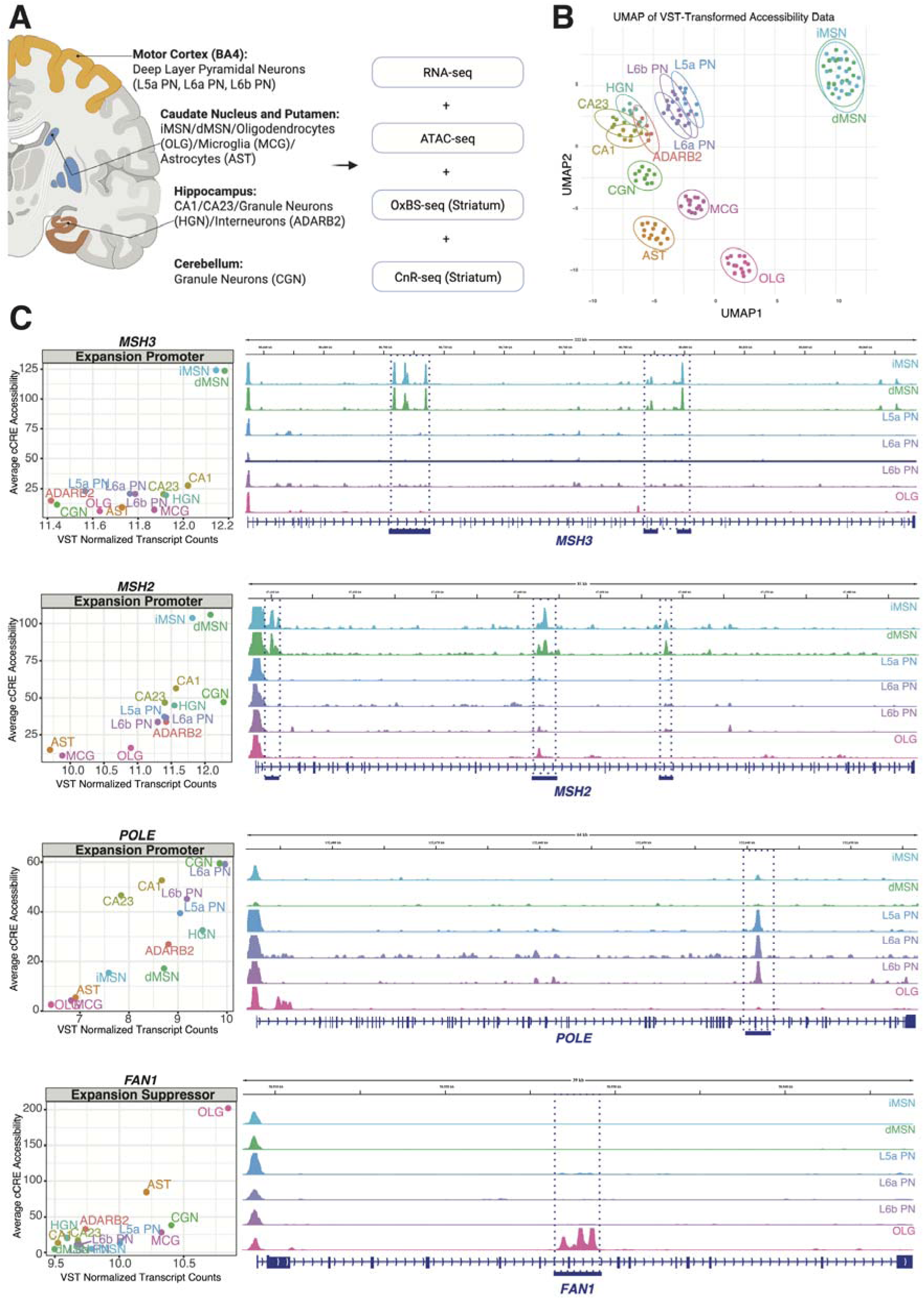
Cell type specific enhancer elements in genes modifying CAG repeat expansion. **(A)** Schematic of the experimental workflow. Nuclei from the following cell types were collected for RNA-seq and ATAC-seq profiling: L5a, L6a, and L6b projection neurons (motor cortex, BA4); direct and indirect medium spiny neurons (dMSNs and iMSNs), oligodendrocytes, microglia, and astrocytes (caudate and putamen); CA1 and CA2/3 projection neurons, granule neurons, and ADARB2 interneurons (hippocampus); and granule neurons (cerebellum). In-depth profiling of striatal MSNs and glia included CUT&RUN-seq for histone modifications, as well as OXBS-seq analysis of CpG and CpH methylation and hydroxymethylation. Created with BioRender.com. **(B)** UMAP dimensional reduction of top 5000 variable accessible regions across all cell types reveals clustering of consensus ATAC-seq datasets by both cell type and brain region. **(C)** Cell type-specific (+)cCRE accessibility in genes modifying of HD onset or progression (*MSH3, FAN1)* or somatic CAG repeat expansion in mouse models *(MSH2, MSH3, POLE, FAN1).* (Left) Normalized ATAC-seq read counts in all (+)cCREs were averaged across control donor samples by cell type and plotted against the average VST-transformed read counts of FANS-seq datasets collected for the respective cell types from control donors’ samples. (Right) Representative IGV tracks showing selected (+)cCREs (blue bars) linked to *MSH3, MSH2, POLE, and FAN1*.

To construct accurate regulatory models, we performed peak-to-gene linkages by integrating RNA-seq expression data with ATAC-seq consensus accessible regions. Using this approach (fig S1A), we functionally annotated regions as either activating (+) or repressive (–)cCREs (table S1). We identified cell-type-specific (+)cCREs associated with genes previously linked to HD onset or progression in GWAS and with components of the mismatch-repair complex influencing somatic CAG expansion in mouse and human cell models (Fig 1C, fig S2a, S3b,c; table S1) ^6,8–11,25,26^. For example, distinct (+)cCREs were observed in intron 7 of *MSH2* and between exons 8–9 of *MSH3* in MSNs, between exons 40–41 of *POLE* in deep-layer cortical neurons, and between exons 7–9 of *FAN1* in astrocytes and oligodendrocytes (Fig. 1c, S2b). Comparative analyses showed that the *MSH3* (+)cCREs are absent in murine MSNs, indicating that these regulatory sites are human-specific (fig S3b,c). Furthermore, cell specific (+)cCREs in MMR genes display both H3K27ac enrichment (fig. S2a,b) and CpG demethylation (fig S2c), confirming their identity as active transcriptional enhancers. In contrast, the promoter regions of *MSH2*, *MSH3*, and *FAN1* showed no cell-type-specific differences in accessibility. Taken together, these data indicate that enhancer-driven transcriptional regulation plays an important role in the cell type specific expression of genes that determine the propensity for somatic expansion in the human brain.

### Decreased DNA demethylation in a subset of enhancers drives transcriptional repression

Studies in mouse models and human HD tissue have consistently shown that cell types undergoing somatic CAG expansion exhibit the most severe transcriptomic alterations ^3–5,9^ (fig S4c). To understand the extent of chromatin alterations that accompany transcriptional dysregulation, we used cell type specific comparative analysis of nuclei from control individuals and those with HD to map chromatin accessibility and cytosine modifications. The magnitude of chromatin dysregulation closely correlated with the mean somatic length gain of the *HTT* CAG tract (Fig 2A, fig S4b), consistent with the observation that the number of differentially expressed genes is higher in cell types with greater CAG repeat instability (fig S4b,c). Furthermore, the chromatin response to *mHTT* somatic CAG expansion was strikingly cell type specific: of the peaks mapped in each cell type, the maximum responses were still a small fraction (∼15%) of the total (Fig 2A) and there was very little overlap between cell types (Fig 2B). Although approximately half of the differentially accessible regions overlapped between striatal dMSNs and iMSNs, the fraction of differentially accessible regions shared between deep layer cortical neurons dropped to less than 20% and there were essentially no shared differentially accessible regions between the striatal and cortical neurons (Fig 2B).

**Fig 2.**
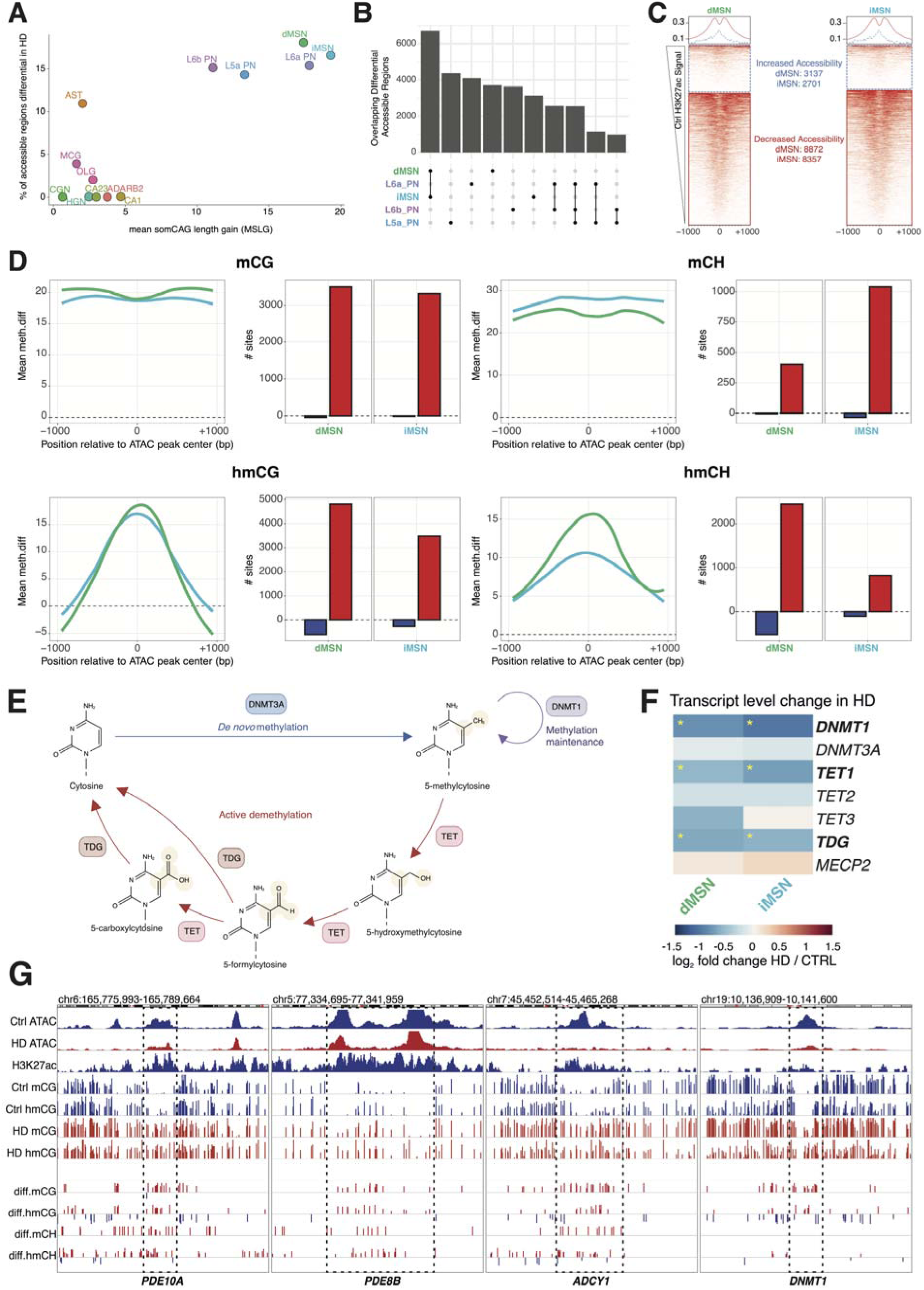
Somatic *mHTT* CAG expansion drives accessibility loss and DNA demethylation deficits at a subset of cell type specific enhancers. **(A)** The relationship between cell type-specific somatic *mHTT* CAG instability (Mean Somatic Length Gain) and percent of accessible chromatin regions that undergo accessibility change in HD (padj < 0.05). **(B)** overlap of differentially accessible regions from (a) in cell types that undergo the most dramatic CAG expansion. **c,** Normalized H3K27ac signal ((0-1, white and red, respectively) in iMSN and dMSNs over regions from (a) separated into those that increase and decrease in accessibility in HD (log2FoldChange > 0, or < 0). **(D) (left),** Metapeak plots centered on decreased accessible regions from (c) with flanks extending +/- 1000kb (80bp bins), with the mean HD-associated change in (hydroxy)methylation (meth.diff) across significant (qvalue < 0.05) cytosines separated into CG and nonCG contexts **(D) (right),** the total number of cytosines in decreasing (+)cCREs that either increase (red) or decrease (blue) significantly in (hydroxy)methylation. **(E)** Schematic of DNA de novo, maintenance and demethylation pathway in post mitotic neurons, created with Biorender.com **(F)** Heatmap depicting disease-associated changes in transcript levels of selected genes regulating DNA methylation. Statistically significant differences are marked with an asterisk (padj < 0.05 by DESeq2, adjusted for multiple comparisons) **(G)** Representative IGV tracks from dMSNs showing decreased accessibility over H3K27ac-positive enhancers linked to *PDE10A*, *PDE8B*, *ADCY1* and *DNMT1*, with differential tracks displaying the meth.diff at individual significant (qvalue < 0.05) CG or CH dinucleotides

Cell types with high CAG instability displayed a bias toward cCREs that lose accessibility (fig S5a). In striatal MSNs, for example, H3K27ac is enriched in ∼8,500 cCREs containing central nucleosome free regions which decrease in accessibility in HD, (Fig 2C) supporting their status as active enhancers ^27^. Decreased accessibility of these enhancers is correlated with decreases in expression of their cognate genes (fig S5b), indicating that inhibition of enhancer function is an important feature of MSN pathogenesis. Most of the ∼3000 cCREs peaks with increased accessibility in HD MSNs are depleted of H3K27ac and are not preferentially associated with increased expression in HD MSNs suggesting that they are not active enhancers (fig S5b).

Differentiated cell types have been reported to require active TET mediated DNA demethylation at a subset of active enhancers ^28,29^ and DNA demethylation is required for proper function of postmitotic neurons ^30^ (Fig 2E). Therefore, we next examined DNA methylation and hydroxymethylation in the repressed enhancer subset. Although accumulation of mCG and mCH occurs across the enhancer domain in HD MSNs, increases in 5hmCG and 5hmCH are localized in the center of the peaks overlapping with the nucleosome depleted regions that are thought to be sites of transcription factor complex interactions (Fig 2D). These enhancers map to previously identified genes strongly repressed in HD dMSNs and iMSNs (*PDE10A, PDE8B, CNR1* and *DNMT1*) ^5^ (Fig 2G, S5b). Furthermore, *TET1* and *TDG*, which mediate sequential steps of active DNA demethylation ^31,32^, are both significantly repressed in HD MSNs (Fig 2F). These data suggest strongly that the accumulation of 5hmCG and 5hmCH, which are intermediates in the DNA demethylation pathway (Fig 2E) ^32^, reflects reduced activity of this pathway in HD MSNs. Since the presence of either 5mC or 5hmC can inhibit the binding of a wide range of site-specific transcription factors ^30,32,33^, we conclude that that their accumulation plays an important role in transcriptional repression in HD MSNs.

### Altered gene body chromatin structure and DNA methylation/hydroxymethylation states reveal two additional epigenetic mechanisms driving transcriptional toxicity in HD

Although the enhancer mediated repression established above occurs specifically in a subset of genes with decreased expression in HD MSNs, it does not explain the entire set of genes whose expression is decreased in HD MSNs or those genes whose expression increases. To identify additional regulatory events that contribute to HD pathogenesis, we analyzed genome-wide transcriptional, histone acetylation, and DNA methylation data in control and HD MSNs (fig S5c). Unbiased clustering of baseline and HD-associated changes in methylation resulted in the definition of three distinct regulatory classes altered during HD pathogenesis (Fig 3A,B, S6a).

**Fig 3.**
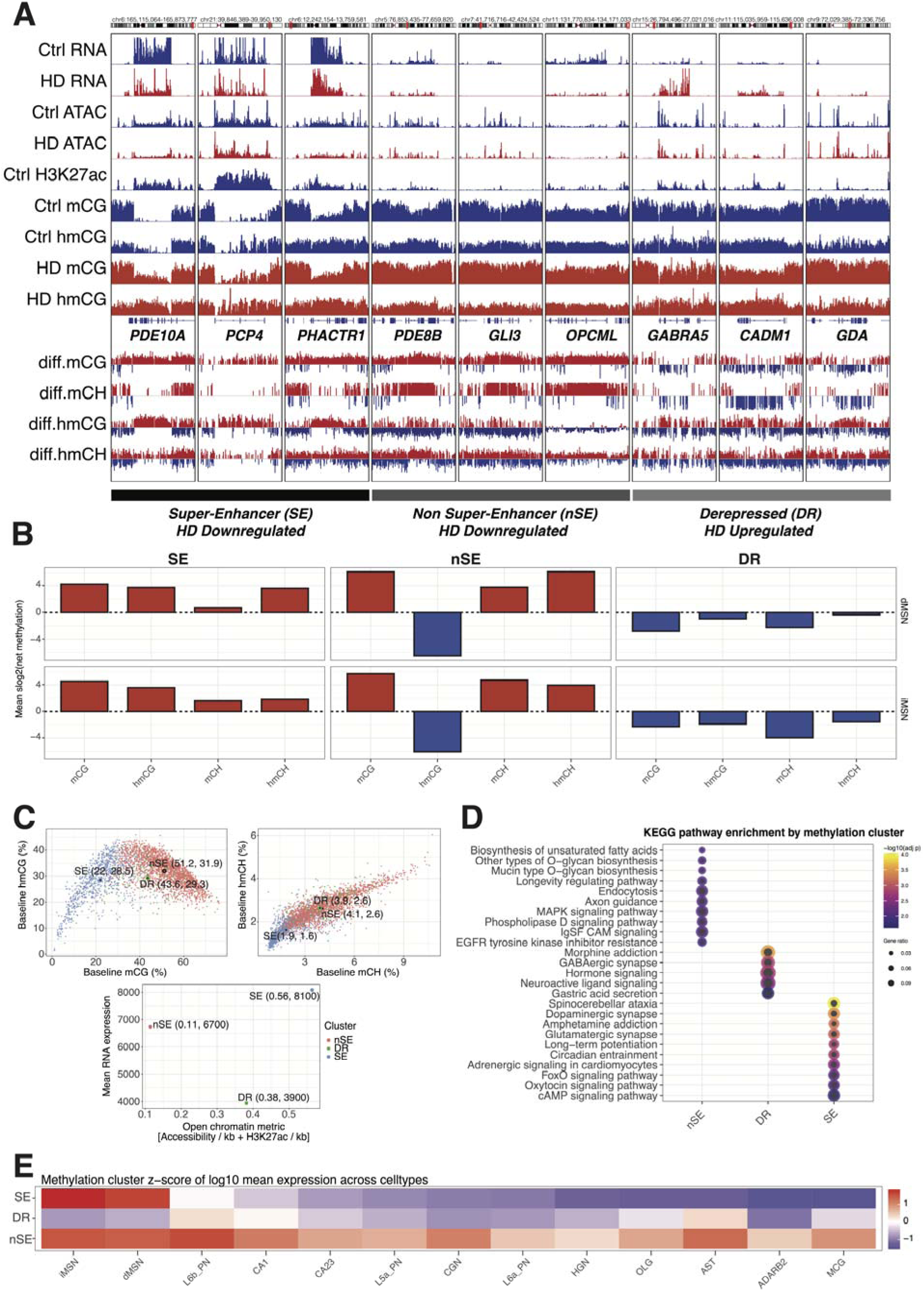
Transcriptional responses to *mHTT* arise from locus specific chromatin states, transcription rate, and unique cytosine methylation dynamics. **(A)** Representative IGV tracks showing transcript level, accessibility, H3K27ac, CpG methylation and hydroxymethylation, and differential methylation across examples of three different classes of genes defined by methylation in control donor MSNs, and change in HD: Super Enhancer (SE) downregulated (*PDE10A, PCP4 and PHACTR1*), Non Super Enhancer (nSE) downregulated (*PDE8B, GLI3, OPCML*), and Methylation Dependent derepressed (DR) (*GABRA5, CADM1, GDA*). Data from dMSNs is shown. **(B)** The average net change in the number of sites that increase and decrease significantly in mCG, hmCG, mCH or hmCH per gene within each class represented as a signed log2(number of sites up minus down) **(C) (top),** (Hydroxy)methylation values are represented as average (%) values of all cytosines in that context and modification, with the relationship between control values for mCG vs hmCG and mCH vs hmCH displayed, the centroid for each cluster a defined value as coordinates is displayed. **(C) (bottom),** Derived metric for ‘openess’ of a given gene is displayed as the sum of the control log10(H3K27ac) and log10(Accessibility) normalized to gene length in kb vs the control expression level of that gene, the centroid for each cluster a defined value as coordinates is displayed. **(D)** KEGG pathway enrichment analysis for all genes in each cluster with the top 10 terms per cluster displayed with padj < 0.05 (top 5 for DR, as <10 terms were under the significance cutoff) **(E)** Global z-score of mean log10(expression) of all genes in each cluster across the cell types profiled in this study, arranged form left to right by decreasing SE values.

First, genes regulated by super enhancer (SE) and repressed in HD MSNs such as *PDE10A, PCP4*, and *PHACTR1* are associated with clusters of co-regulated enhancers, broad H3K27ac enriched regions over the gene body, and near-complete gene-body demethylation and are highly expressed in MSNs compared to other cell types (Fig. 3A,C,E fig S6a). These properties are typical of SE regulated genes that define differentiated cell types ^34^. In HD MSNs SE genes primarily gain mCG and hmCG in their gene bodies as well as their enhancers (Fig 3A,B). These data are consistent with decreased DNA demethylation as the driver of their dysregulation in both of these domains. A second non-super enhancer regulated (nSE) subset of genes (e.g. *PDE8B, GLI3*, and *OPCML*) are also primarily repressed in HD MSNs. These genes are more moderately expressed, are more methylated, and have no significant broad regions of H3K27ac in their gene body (e.g *PDE8B*). Furthermore, nSE genes show concordant increases in mCG, mCH, and hmCH but a pronounced loss of hmCG (Fig 3A,C,E fig S6a) in the gene body. A third subset of genes in MSNs increase in expression HD MSNs, they decrease in 5mCG, 5hmCH, and 5hmCH in the gene body and increase in 5hmCG (Fig 3A,B). These genes, including *GABRA5*, *MAN1A1*, *GPR176*, and *OPRK1*, are moderately expressed and there is no apparent change in chromatin accessibility or DNA methylation status in their associated cCREs (Fig 3C,E). We note that genes in the de-repressed (DR) subset in HD MSNs are associated with DNA methylation loss within the gene body. *DNMT1*, an nSE-repressed gene in HD MSNs, is approximately two-fold downregulated in HD MSNs suggesting that its loss may contribute to the increased gene body DNA methylation evident in DR genes (Fig S5b).

Strong evidence that all three of these dysregulated gene classes contribute to loss of MSN function is provided by KEGG pathway enrichment analysis showing signatures relevant to autophagy regulation in the nSE cluster, inhibitory gPCR signaling in the DR cluster, and cAMP signaling in SE cluster (Fig 3d). Moreover, human genetic studies indicate that individual genes in each of these classes are important for MSN viability or neuronal function in humans (fig S6c, table S2). For example, the SE repressed gene PDE10A (>2X decreased expression) causes Autosomal Dominant Striatal Degeneration 2 (ADSD2), the nSE repressed gene PDE8B (∼2X decreased expression) causes Autosomal Dominant Striatal Degeneration 1 (ADSD1), and gain of function mutations in the DR gene GABRA5 (>2X increased expression) cause Developmental and Epileptic Encephalopathy 79 (DEE79) ^35–37^. There are many other genes identified as strongly dysregulated in figure S5b for which human genetic studies provide evidence of an essential function in the brain.

### The regulation of gene expression is disturbed in the majority of HD MSNs

Quantitative measures of DNA methylation indicate that in many genes from the clusters identified (Fig 3A,B), changes in methylation at specific CG dinucleotides occur in 50-90% of MSN genomes (Fig 4A). This suggests that epigenetic dysregulation and consequent transcriptional repression occurs in a large fraction of HD MSNs. To directly measure the impact of these events on individual HD MSNs, we used RNAscope to measure changes in the mRNA abundance of *PDE10A*, *PCP4,* and *CALM2* three genes whose expression change by more than 50% in HD, relative to the presence of an MSN-specific mRNA whose expression does not significantly change in HD (*COCH*) (*5*). In sections from control donor samples, *PDE10A*, *PCP4* and *CALM2* (Fig 4B-D) were detectable in the majority of *COCH-*positive MSNs across donors. As expected, although the number of MSN nuclei were substantially reduced in HD donor samples, *COCH* mRNA was present at comparable levels in the remaining MSNs, whereas very low levels of *PDE10A*, *PCP4,* and *CALM2* (Fig 4B-D) were detected in these same neurons. Quantitative image analysis demonstrated definitively that reduced expression of these genes occurred i n the majority of HD MSNs (Fig 4E-G). Taken together, changes in DNA methylation revealed by oxBSseq (Fig 4A), and mRNA abundance by RNAscope (Fig 4E-G) demonstrate that most striatal MSNs experience marked transcriptional dysregulation and cellular dysfunction in HD patients.

**Fig 4.**
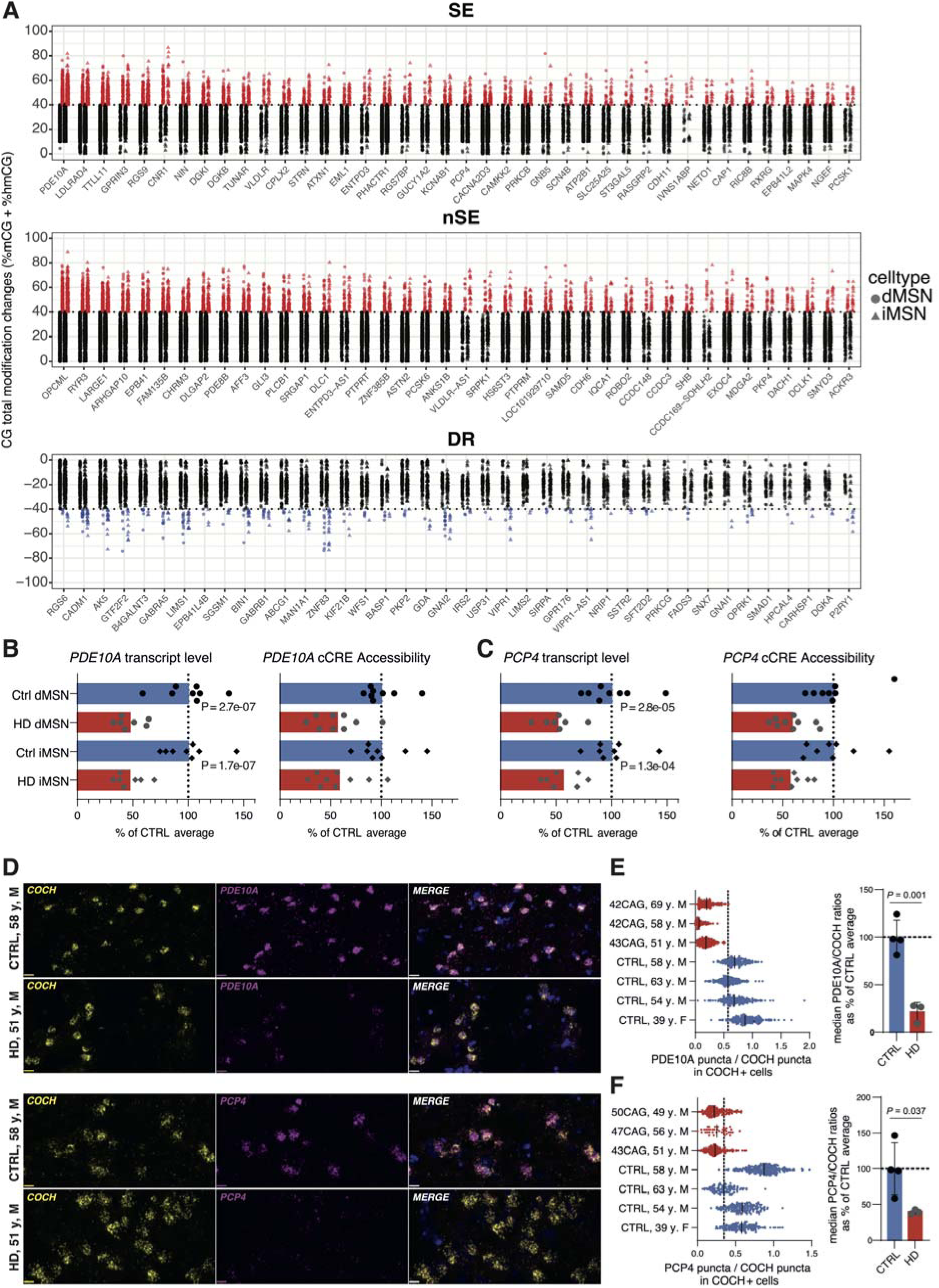
Gene expression is disrupted in the majority of MSNs in HD. **(A)** Total significant (qvalue < 0.05) CG methylation change (hmCG_meth.diff + mCG_meth.diff) in the top 40 genes from the each of the clusters identified in figure 3 with the maximal HD associated methylation change. Individual sites increasing more than 40% in repressed (log2FoldChange < 0) genes from SE and nSE and decreasing more than 40% in derepressed (log2FoldChange > 0) genes from DR are highlighted in red and blue, respectively. **(B, C)** *PDE10A* and *PCP4* transcript level and cCRE accessibility in HD MSNs represented as % of their transcript level and accessibility in control donor MSNs (mean plotted). pvalues were calculated with DESeq2 and are adjusted for multiple comparisons. **(D)** Representative sections of putamen stained with RNAScope for the quantification of *COCH* (yellow) and, *PDE10A* and *PCP4* (magenta) transcripts (cell nuclei stained with DAPI, blue). **(E,F)** Ratio of *PDE10A*+ puncta to *COCH*+ puncta and *PCP4*+ puncta to *COCH*+ puncta was calculated for each *COCH*+ cell (median of the ratio is plotted for cells from each tissue donor analyzed). Vertical dashed line indicates the lowest median *PDE10A/COCH* and *PCP4/COCH*, ratio out of the four control donors analyzed. The average of median *PDE10A/COCH* and *PCP4/COCH* ratios in HD donors was significantly lower than in control donors (P = 0.001, P = 0.0368, for *PDE10A*, *PCP4* respectively, using unpaired two-tailed T-test, average +/- SD is shown).

## Discussion

To understand selective cellular vulnerability and identify therapeutic opportunities in late onset neurodegenerative disease detailed knowledge of molecular events accompanying pathogenesis in the human brain is essential. Here we have used high resolution epigenetic profiling of human postmortem samples to reveal a complex and previously unrecognized pathogenic cascade in Huntington’s disease. The data presented here clarify the mechanism that establishes regional and cell type specific expression of genes that have been associated with somatic instability in the initial stage of HD and establish DNA demethylation as a critical factor in the second, toxic phase of pathogenesis. Therefore, our findings provide a framework for understanding the impact of expanded mutant huntingtin on the functions of MED15 and TCERG1 in HD pathogenesis ^38,39^, and they identify new therapeutic targets that may allow safer long-term treatment and improved efficacy for patients suffering from this devastating disease.

An important first insight into HD pathogenesis reported here is that the regional and cellular specificity of somatic expansion in the human brain is driven by cell type, and in some cases species, specific transcriptional enhancers in mismatch repair (MMR) genes. The data predict that the presence of enhancers in *MSH2* and *MSH3* in MSNs, and the activity of the *FAN1* enhancer specifically in glia, contribute to the differential expression of these genes, resulting in a high MutSβ to FAN1 ratio ^5,9,10^, thus promoting *mHTT* CAG expansion. The inverse situation in astrocytes and oligodendrocytes suggests that these enhancers contribute to *mHTT* CAG stability. Given the ongoing therapeutic initiatives aimed at inhibition of somatic expansion and the critical roles of MMR proteins MSH2, MSH3, POLE and FAN1 in repeat instability ^5,6,8–12,25,26,38–40^, we propose that the human MMR gene enhancers present new opportunities for therapeutic intervention in HD. This strategy may reduce the propensity of somatic expansion in critical brain structures while at the same time preserving DNA repair functions required for genome stability in the CNS and periphery.

A second advance we report here is the delineation of a complex cascade of epigenetic events that drives toxicity in HD. Integration of the high definition ATACseq, H3K27ac mapping and genome wide, single nucleotide resolution DNA methylation data presented here with results from animal models ^5,15,18–20,41^ and human genetic studies ^6–8,14^ describe a succession of epigenetic events that support a new mechanistic hypothesis of HD pathogenesis (Fig. 5). In the first step, as discussed above, the specificity of somatic expansion and selective cellular vulnerability in the human brain results from enhancer mediated transcription of MMR genes that promote or oppose expansion in *mHTT* exon1. A second step then ensues as a result of emHTT interactions with HD-GWAS ^6,8^ age of onset modifiers MED15 ^38^ and TCERG1 ^41^ that disrupts Mediator complex functions at enhancers and the efficiency of transcription in the gene body for both SE and nSE gene clusters. This results in transcriptional dysregulation of thousands of genes, including *TET1, TDG* and *DNMT1*. This reduced DNA demethylation further compromises expression of SE and nSE genes, which include *DNMT1,* could contribute to derepression in yet a third class of genes. In these DR genes, decreased gene body DNA methylation results in diminished binding of neuronal repressor proteins such as MeCP2 ^42,43^, elevating their expression. Despite the complexity of these concurrent and overlapping epigenetic events, the simplest interpretation of the data is that decreased DNA demethylation is critical in driving toxicity and progression in HD. We propose, therefore, that development of therapeutic strategies to reverse toxicity in HD should incorporate efforts to elevate TET activity in neurons carrying expanded mutant huntingtin.

**Fig 5.**
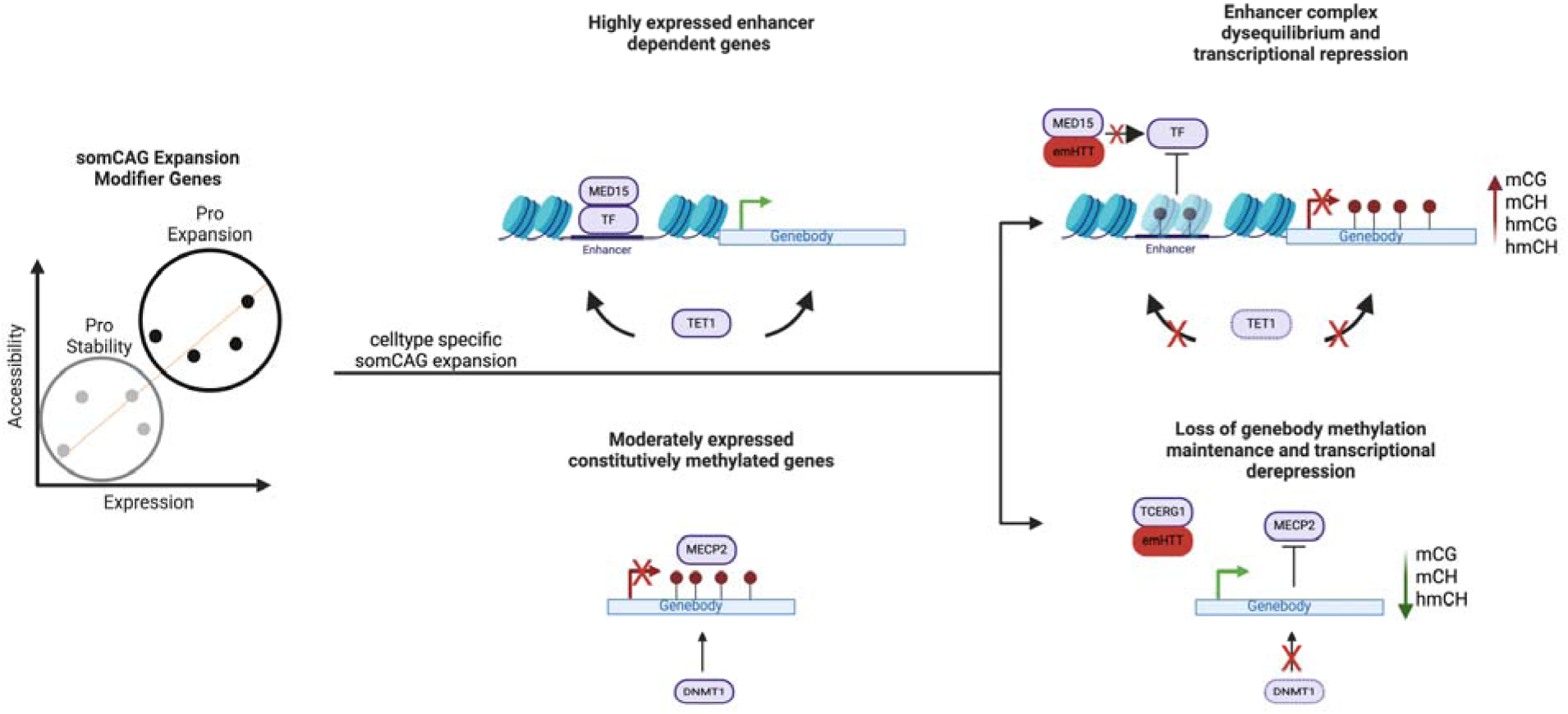
Model of human Huntington’s Disease pathology in MSNs. Phase 1: Celltype specific accessibility of enhancers in genes encoding proteins involved in MMR drive high expression of genes encoding complexes such as MutSB (MSH2/MSH3) which promote somatic CAG expansion of *mHTT,* or FAN1 which promotes CAG stability Phase 2: In cell types that undergo somatic CAG expansion, the resulting emHTT (protein), which has been shown to interact with MED15 and TCERG1 (38, 39), disrupts enhancer complex formation and DNA demethylation equilibrium leading to transcriptional repression of a subset of enhancer dependent genes. Concurrently, DNA methylation maintenance is lost, and aberrant transcription increases, at a subset of moderate to lowly expressed genes leading to their derepression.

Finally, we note that there are specific functionally important sites where, in HD, CpGs have gained or lost methylation and hydroxymethylation in a majority of MSN genomes. Additionally, genes associated with those repressed sites, such as *PDE10A* and *PCP4*, are similarly repressed in >50% of nuclei via RNAscope validation of mRNA abundance. This demonstrates that transcriptional toxicity occurs in majority of HD MSNs, including genes known to cause human disease such as autosomal dominant striatal degeneration 2 (PDE10A, SE class) ^37^ or haplo-insufficient adult-onset striatal degeneration 1 (PDE8B, nSE class) ^35,44^ and other CNS disorders ^45–48^. These data suggest that these mechanisms result in significant dysfunction in HD MSNs. We conclude, therefore, that in patients already experiencing symptoms, treatments that combine inhibition of somatic CAG expansion with TET mediated reversal of DNA demethylation deficiencies will be essential for maximal efficacy.

## Acknowledgements

We are thankful to T.F. Vogt, V. Beaumont and J. Chen for helpful discussions throughout the project. We thank Karen Liu and C. Wang for technical assistance. We are also grateful to the Rockefeller University Genomics Resource Center for advice and support. We thank J.P.G. Vonsattel and A.F. Teich from Columbia University Alzheimer’s Disease Research Center, D. Keene from University of Washington BioRepository and Integrated Neuropathology Laboratory, S. Berretta and the Harvard Brain Tissue Resource Center, The University of Michigan Brain Bank, University of Maryland Brain and Tissue Bank, Netherlands Brain Bank, and UCLA Human Brain & Spinal Fluid Resource Center for assisting and providing post-mortem brain tissues.

## Funding

CHDI Foundation (MB, KM, NH, II, HC, ES, YH, LK, CP, ND, ES)

## Author contributions

Conceptualization: M.B., K.M., N.H. Formal analysis: M.B., K.M., I.I., M.E.S., H.C., T.S.C., D.B., C.P., N.D. Investigation: M.B., K.M., I.I., M.E.S., Y.H., E.S., P.D., C.P. Resources: L.K., N.D. Data Curation: M.B., L.K., N.D. Visualization: M.B., K.M. Writing of original draft: M.B., N.H., K.M. Writing, review and editing: K.M., M.E.S., I.I., Supervision: N.H. Funding acquisition: N.H.

## Competing interests

The authors declare no competing interests.

## Data and materials availability

All sequencing datasets generated as part of this study will be publicly available in NCBI GEO and are available for the review process with the provided token access key (GSE312285 (token:wdsrauwcndipfuj), GSE312283 (token: gxqpkucyzburnmp), GSE312279 (token: yhqjsumwtvwpfol), GSE312277 (token: ovwtuakqvnyhpsb)). Further information and requests for resources and reagents should be directed to the lead contact, N. Heintz. The publicly available resources used (reference genome sequence and annotation, and computational analysis tools) are described in the Methods section. Only publicly available tools were used in data analysis. The analysis parameters used have been described in Methods.

## Supplementary Materials

### Supplemental Figures S1-S7, Supplemental Note 1

**Fig S1.**
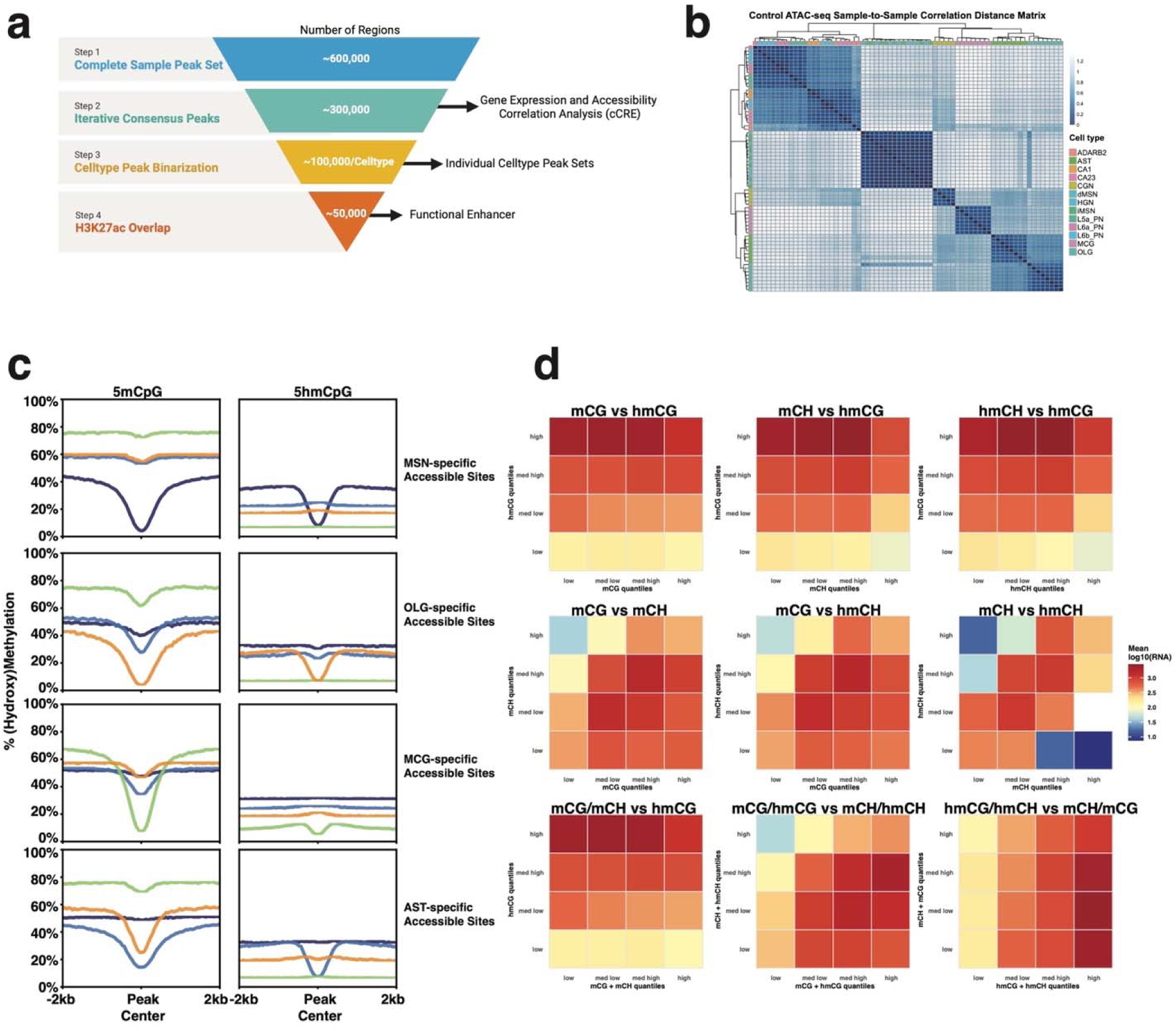
ATACseq processing, and the relationship of methylation to accessibility and expression. **a,** The analysis of chromatin accessibility data included (step 1) peak calling, (step 2) identification of robust and reproducible consensus peaks and, as a subset, cCREs that correlate positively or negatively with transcript levels in matched FANS-seq datasets, (step 3) peak binarization-based definition of peak sets for each of the cell types, and their use in the analysis of HD-associated differential accessibility and (step 4) integration with H3K27ac CUT&RUN-seq datasets to identify accessible chromatin regions that are active enhancers. **b,** Distance clustering matrix of chromatin accessibility profiles of individual cell type-specific ATAC-seq datasets from control donors, based on normalized counts over iterative consensus peaks. **c,** The relationship in dMSNs between genebody mCG, hmCG, mCH and hmCH, the sum of modifications in CG and nonCG contexts, sum of hydroxymethylation in CG and non CG contexts, and sum of methylation in CG and nonCG contexts, and their effects on expression displayed as quartiles (low = bottom 25%, med low = 25-50%, med high = 50-75%, high = top 75%)

**Fig S2.**
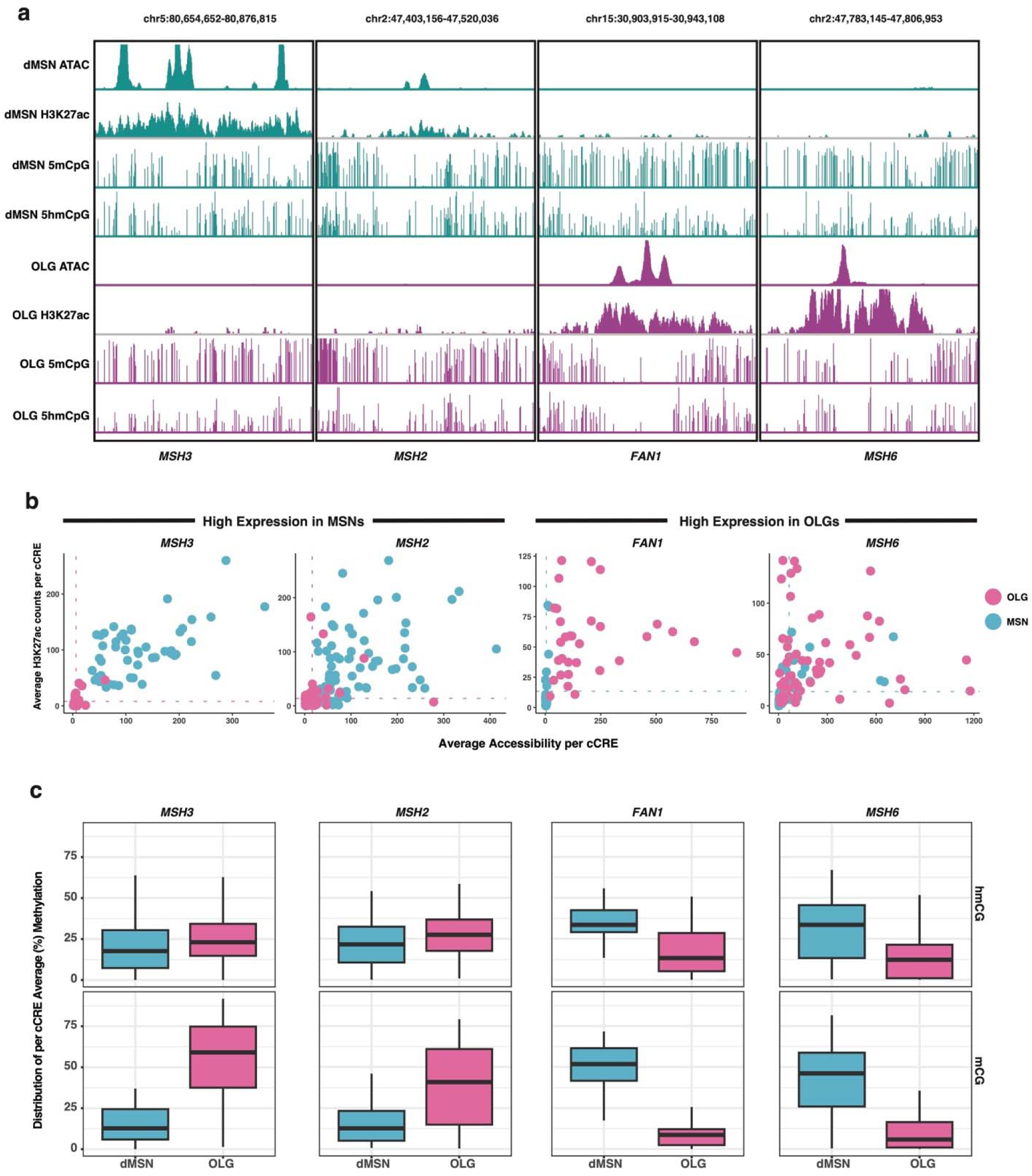
cCRE enhancer validation via DNAme, accessibility, and H3K27ac. **a,** Representative IGV tracks showing selected (+)cCREs linked to *MSH3, MSH2, FAN1 and MSH6,* and their accessibility, H3K27ac signal and level of 5mCpG and 5hmCpG in dMSNs and oligodendrocytes. **b,** Normalized counts of ATAC-seq and CUT&RUN-seq reads from dMSN and oligodendrocyte samples were plotted to compare the levels of accessibility and H3K27ac at individual (+)cCREs linked to *MSH2* and *MSH3* (higher expression in MSNs) or to *FAN1* and *MSH6* (higher expression in oligodendrocytes). Dotted lines represent the mean ATAC-seq signal (vertical) or mean H3K27ac signal (horizontal) over all (+)cCREs in the cell type with lower expression of the gene. **c,** Level of CpG hydroxymethylation and methylation over (+)cCREs linked to genes with higher expression in MSNs (*MSH2, MSH3*) or in oligodendrocytes (*FAN1, MSH6*). The mean, Q1 and Q3 of all (+)cCREs are shown.

**Fig S3.**
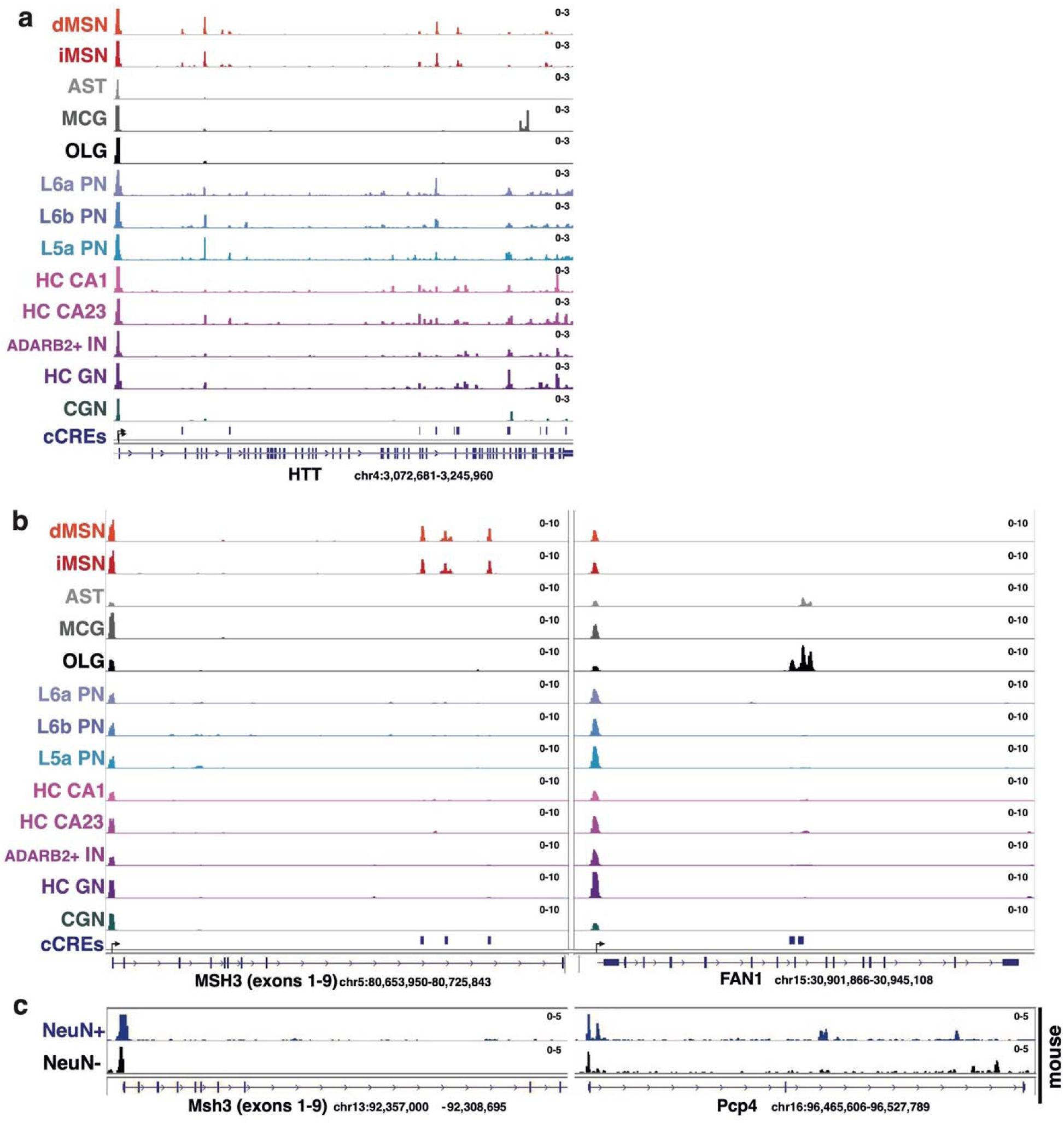
Cell type-specific accessible regions in *MSH3*, *FAN1* and *HTT* genes. **(A,B)**, Representative distribution of sequencing reads over (a) *HTT*, (b) *MSH3* and *FAN1* genes in ATAC-seq datasets generated from nuclei of human brain cell types. The position of cCREs linked to these genes is shown. **(C)** Distribution of sequencing reads over *Msh3* and *Pcp4* genes in ATAC-seq datasets generated from anti-NeuN-positive and -negative nuclei isolated from mouse striatum. Accessibility at neuronal marker genes (like *Pcp4*) was used to assess purity of isolated populations.

**Fig S4.**
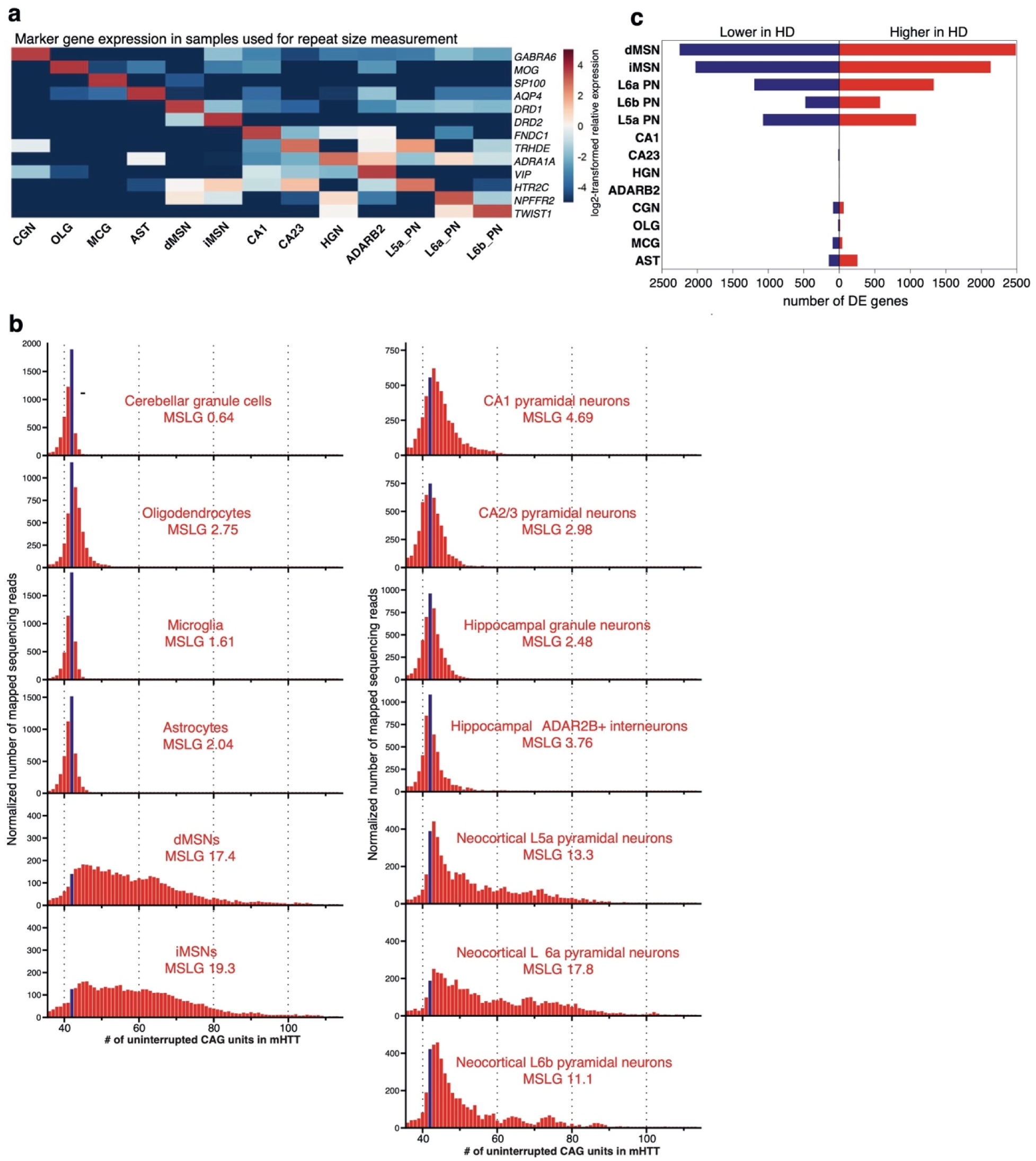
Somatic expansion of *mHTT* and transcriptional dysregulation in the cell types analyzed. **(A)** Relative expression level of marker genes in the transcriptome of nuclei collected for *mHTT* CAG repeat length analysis. Heatmap depicts log_2_-transformed relative expression in each sample (calculated based on DESeq2-normalized counts). **(B)** Length distribution of the *mHTT* CAG repeat in different cell types in two 69-year-old male donors that carried a tract of 42 uninterrupted CAG units. Data from striatal glia is derived from donor 1, the rest of the cell types from donor 2. In cell types analyzed from both donors, there were no notable inter-donor differences (not shown). Blue bar marks sequencing reads derived from the initial unexpanded CAG tract. y axes denote normalized number of sequencing reads mapped to reference sequences with different CAG tract lengths (normalized by scaling to 5,000 reads). Reads derived from the normal *HTT* allele are not shown. Median somatic length gain (MSLG) is calculated as in Mätlik et al, *Nat Genet.*, 2024. **(C)** Number of differentially expressed (DE) genes (Padj < 0.05 by DESeq2, adjusted for multiple comparisons) in the comparison of HD and control donor FANS-seq datasets from dMSNs and iMSNs (n = 7 and n = 8 HD and control datasets, respectively), neocortical L6a PNs (n = 19, n = 25), L6b (n = 22, n = 19) and L5a PNs (n = 17, n = 25), hippocampal CA1 (n = 6, n = 7), CA2/3 (n = 5, n = 6), granule neurons (n = 6, n = 7) and ADARB2+ INs (n = 6, n = 4), cerebellar GNs (n = 9, n = 9), and striatal oligodendrocytes (n = 7, n = 7), microglia (n = 7, n = 7) and astrocytes (n = 7, n = 7).

**Fig S5.**
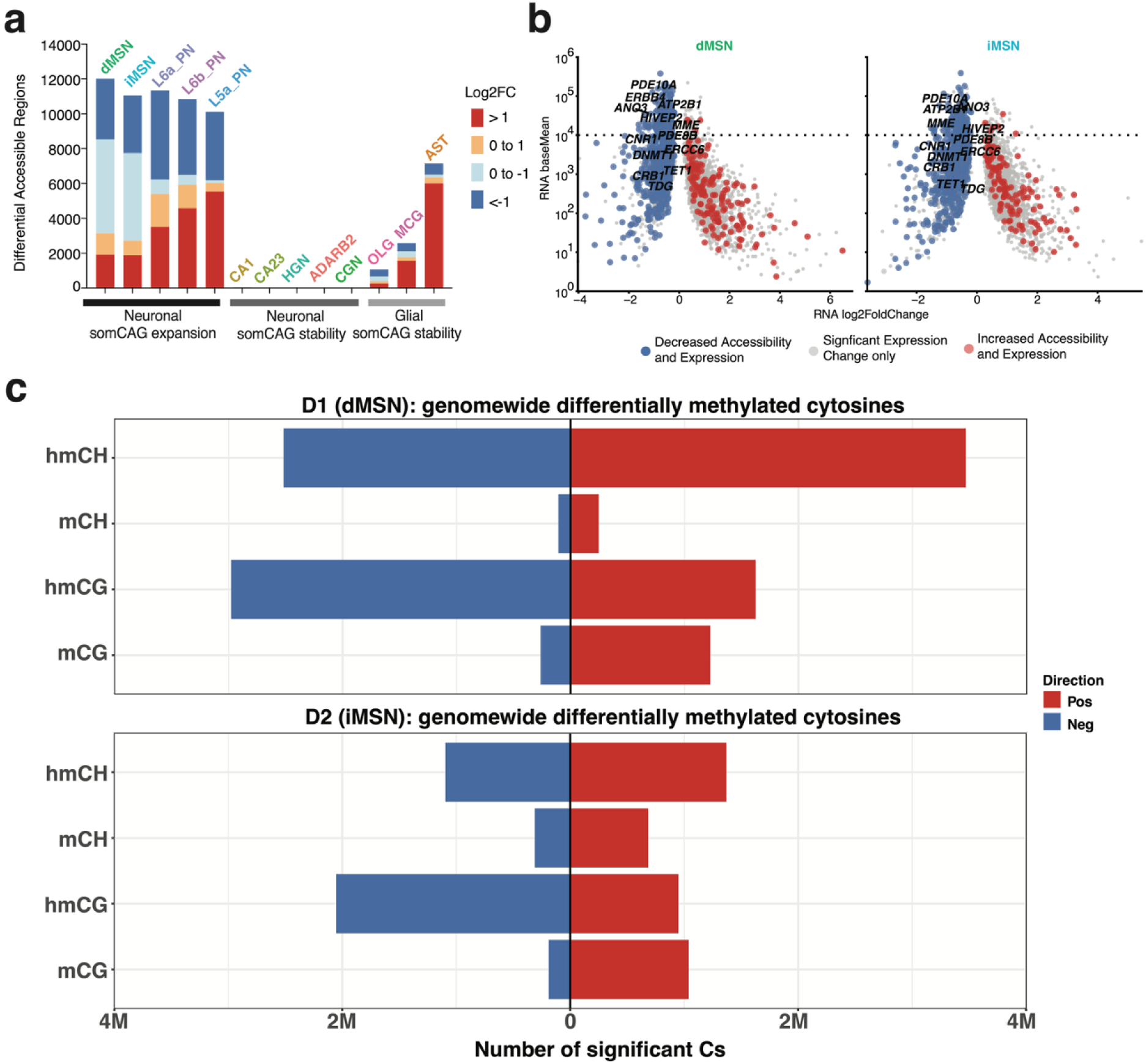
Genome-wide HD associated methylation changes and ATACseq/RNAseq differential. **(A)** The number of differentially accessible chromatin regions (*p adj* < 0.05) in the cell types analyzed. Log2FoldChange < 0 and Log2FoldChange > 0 values indicate reduced and increased chromatin accessibility in samples from individuals with HD, respectively **(B)** Scatter plots of all significant (padj < 0.05) differentially expressed genes in HD dMSNs and iMSNs, marking genes for which repression or derepression correlated with reduced (red) or increased (blue) (+)cCRE accessibility, respectively. Genes that are known to cause haploinsufficient human disease support MSN function (e.g. *CNR1)*, or regulate DNA methylation (*TET1*, *TDG*, *DNMT1*) are labeled. **(C)**The number of cytosines with significant HD-associated increase (pos) or decrease (neg) in (hydroxy)methylation in CG and nonCG (CH) contexts (qvalue < 0.05) shown for dMSNs and iMSNs

**Fig S6.**
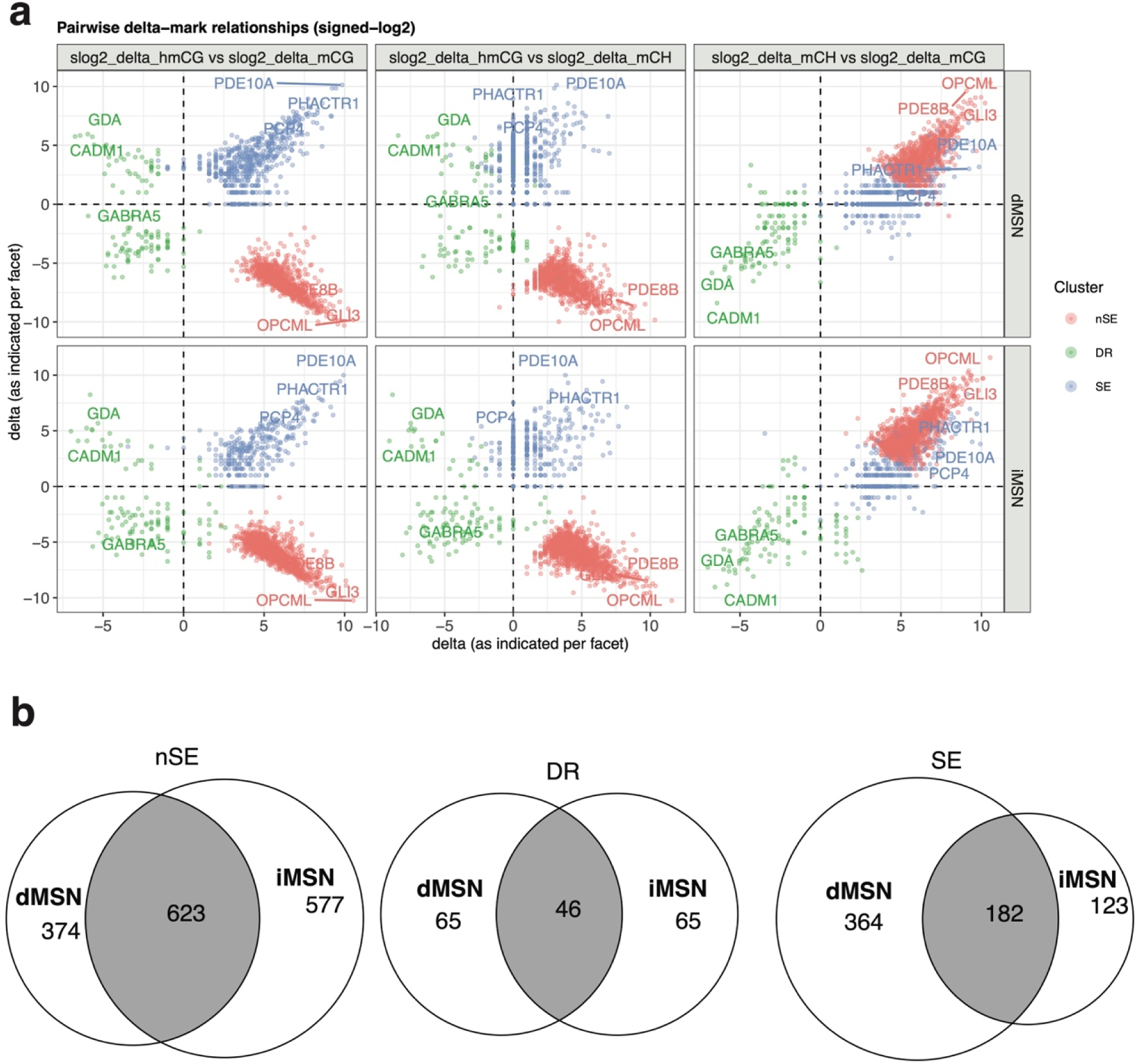
Clustering results of control genic DNAme baselines in dMSN and iMSNs and HD-associated significant cytosine changes in HD differentially expressed genes. For each significantly differentially expressed gene in HD dMSNs or iMSNs, the scatterplots display the *signed log –scaled cumulative change* in cytosine modifications (**slog2(delta)**), calculated as log (1 + the net number of significant cytosines that increase in (hydroxy)methylation minus the number that decrease in (hydroxy)methylation) across mCG, hmCG, mCH, and hmCH sites within the gene body. These values summarize the overall direction disruption per gene. Each point represents a single gene, colored according to unbiased k-means cluster membership defined by slog2(delta) and control baseline genebody methylation (the average % (hydroxy)methylation level across all cytosines per each context) **(B)** the overlap between differentially expressed genes assigned to each cluster in dMSNs and iMSNs.

**Fig S7.**
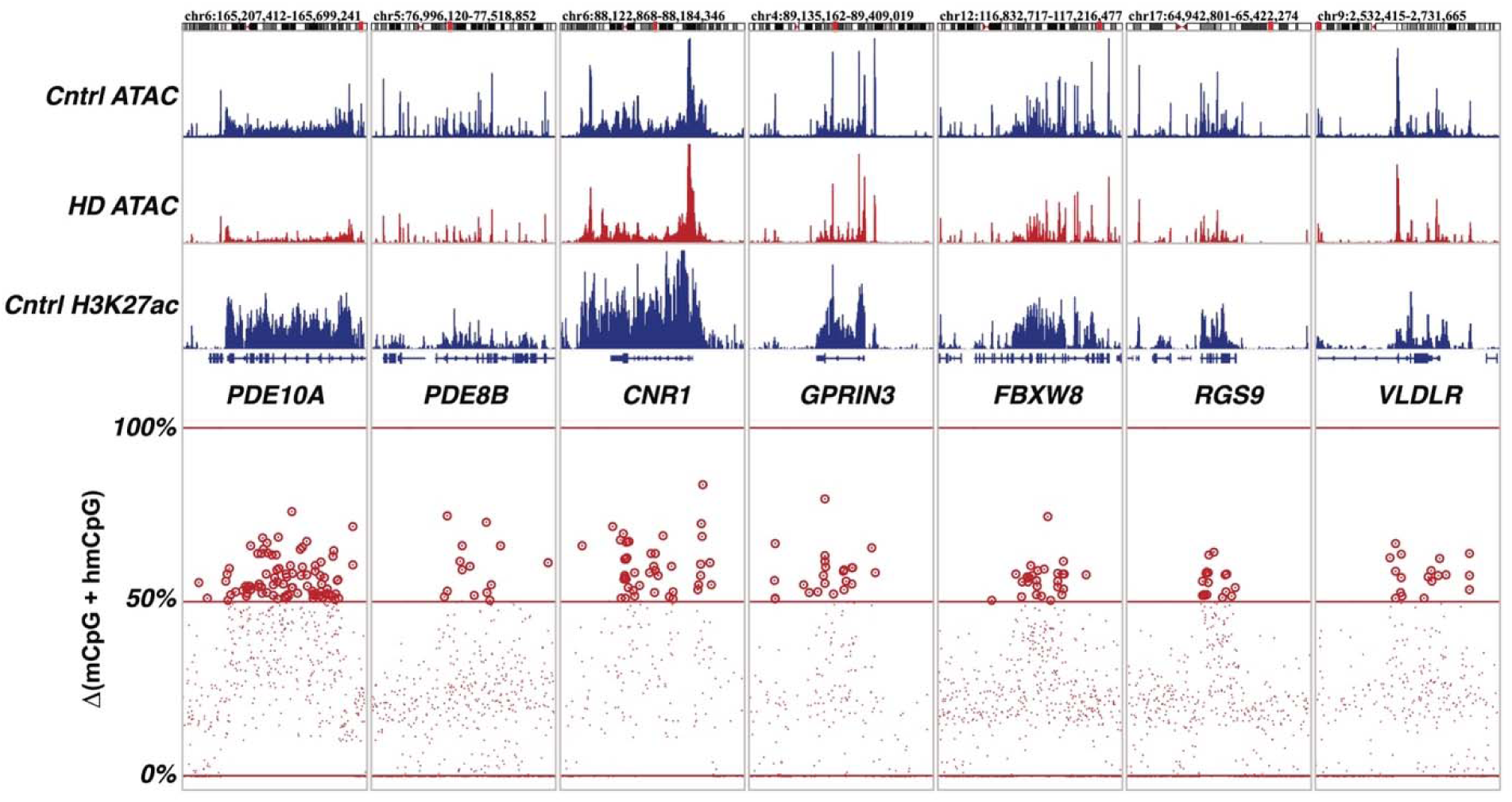
Total change in CG modification level in selected genes with HD associated transcriptional dysregulation. Representative IGV plot showing control and HD dMSN accessibility, Control MSN H3K27ac, and a positional dot plot of the sum of mCG and hmCG meth.diff values for signficant (qvalue < 0.05) cytosines in HD dMSNs (Total change in CG modification level) in genes with decreased expression and accessibility in HD (*PDE10A, PDE8B, CNR1, GPRIN3, FBXW8, RGS9, VLDLR*). Only cytosines with positive meth.diff values are displayed (meth.diff > 0).

**Supplementary Note 1.**
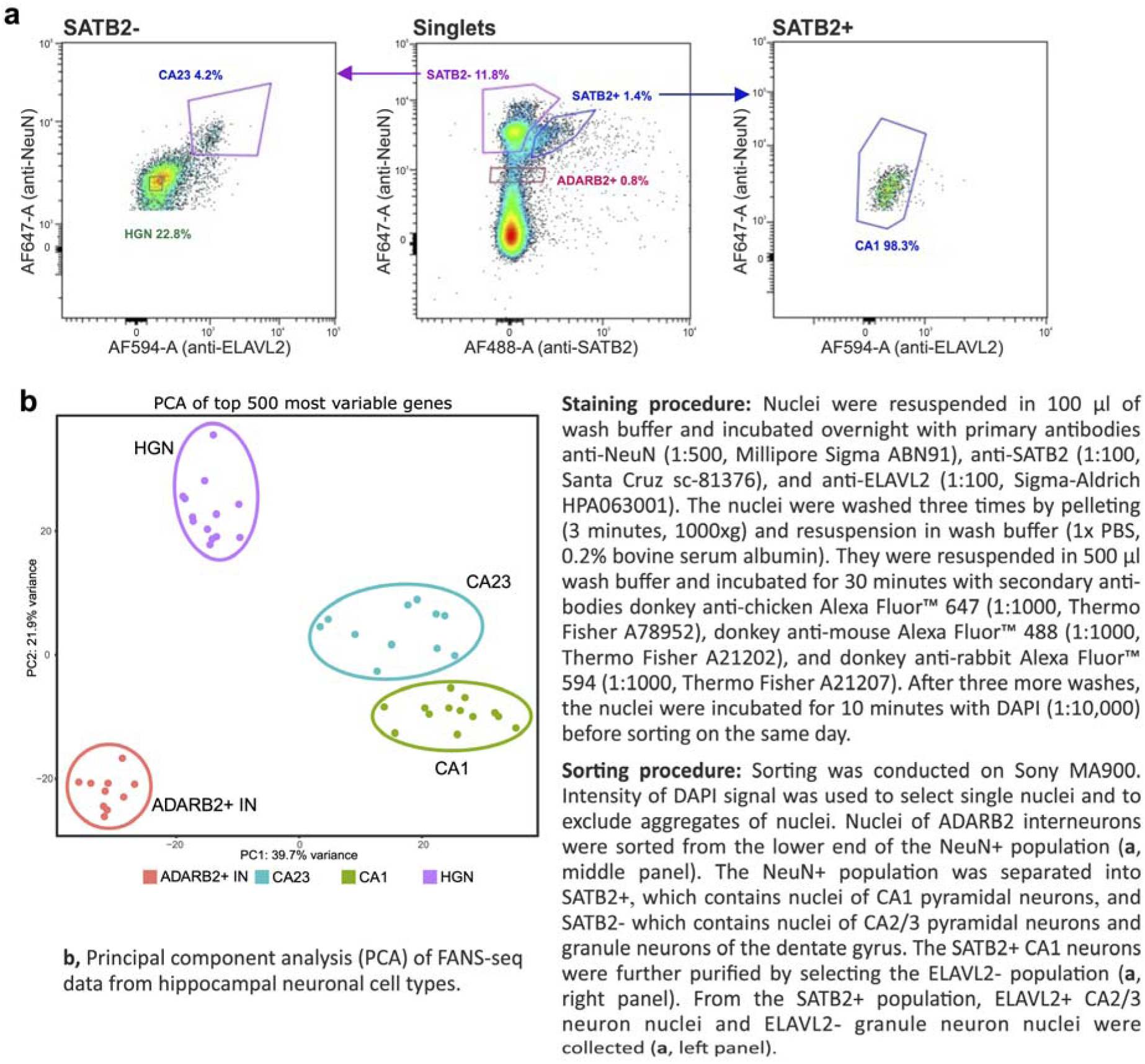
Labeling and sorting of the nuclei of hippocampal neuronal types.

### Methods

#### Human samples

Deidentified tissue samples analyzed in this study were determined to be exempt from Institutional Review Board review according to 45 CFR 46.102 (f). For this work, fresh frozen brain samples were obtained from Miami’s Brain Endowment Bank, University of Washington BioRepository and Integrated Neuropathology Laboratory, Columbia University Alzheimer’ Disease Research Center, University of Maryland, Science Care, and Netherlands Brain Bank or through the National Institutes of Health (NIH) NeuroBioBank and sourced from either the Harvard Brain Tissue Resource Center, The University of Michigan Brain Bank or the NIH Brain & Tissue Repository-California, Human Brain & Spinal Fluid Resource Center, VA West LA Medical Center (Los Angeles, CA). Drug addiction and schizophrenia as well as clinical evidence of brain cancers were reasons for sample exclusion, whereas samples from donors with a history of other non-brain cancers and diabetes were accepted. Caudate nucleus, putamen BA4, hippocampus and cerebellar vermis were used for isolation of nuclei. The brain regions used from each donor and their age, race, sex and post-mortem interval are noted in Supplementary Table 1.

#### Sorting nuclei of striatal MSNs and glia, cerebellar granule neurons, hippocampal neurons and cortical pyramidal neurons

Nuclei were isolated from fresh frozen brain tissue samples as previously described in ***Basic Protocol 1***^21^. Nuclei were labelled and sorted with FANS as previously described in ***Basic Protocol 2*** (neocortical cell types), ***Basic Protocol 3*** (striatal MSNs)^21^, and in ^5^(cerebellar granule neurons and glial cell types of the striatum). The nuclei of hippocampal neuron types were labeled with anti-NeuN (1:500, Millipore Sigma ABN91), anti-SATB2 (1:100, Santa Cruz sc-81376), and anti-ELAVL2 (1:100, Sigma-Aldrich HPA063001) antibodies and with donkey anti-chicken Alexa Fluor™ 647 (1:1000, Thermo Fisher A78952), donkey anti-mouse Alexa Fluor™ 488 (1:1000, Thermo Fisher A21202), and donkey anti-rabbit Alexa Fluor™ 594 (1:1000, Thermo Fisher A21207) secondary antibodies. The nuclei of ADARB2-expressing interneurons (NeuN+^lo^), CA1 pyramidal neurons (NeuN+^hi^, SATB2+ ELAVL2-), CA2/3 pyramidal neurons (NeuN+^hi^, SATB2-, ELAVL2+) and granule neurons of the dentate gyrus (NeuN+^hi^, SATB2- ELAVL2-) were collected as described in **Supplementary Note 1**.

#### Isolation of RNA, gDNA and Tn5-Treated DNA

RNA and gDNA from sorted nuclei, and gDNA from Tn5-treated nuclei were isolated and turned into sequencing libraries as previously described in ***Basic Protocol 4, Support Protocol 1*** and ***Support Protocol 2***^21^. FANS-seq and ATAC-seq sequencing reads were processed and quality-controlled according to ***Basic Protocol 5***^21^.

#### Differential gene expression analysis

Differential gene expression analysis was performed using ‘Genebody.Counts’ (FANS-seq reads mapped to full gene bodies) and DESeq2 ^49,50^ (version 1.40.2). Differential gene expression analysis results were filtered to exclude genes for which none of their annotated TSS positions in NCBI Refseq hg38 (version 109.20211119) overlapped with ATAC-seq consensus peaks defined for the cell types based on presence in presence in at least half of control samples analyzed. For neocortical neuron types and MSNs, previously published lists of differentially expressed genes were used^4,5^. Pheatmap R package (version 1.0.12) was used for visualizing gene expression levels on heatmaps.

#### oxBS-seq Library Preparation and Processing

OxBS-seq libraries were prepared according to manufacturer’s instructions in the Ultralow Methyl-Seq with TrueMethyl^®^ oxBS (Tecan Genomics, #0541-32). Briefly, 100-300 ng of gDNA (∼50,000-150,000 nuclei) with 1% unmethylated lamda phage DNA spike-in was sonicated and purified for 200bp fragments. Purified gDNA fragments were then split into paired samples such that 30 ng was used for BS-seq (mock-oxidation reaction) and the remaining gDNA was used for oxBS-seq (with oxidation reaction). Libraries were indexed, PCR amplified and sequenced to >300million reads per sample on NovaSeq6000 (2 × 100 bp).

The raw OxBS-seq fastq reads were trimmed with the parameters “--paired --fastqc --cores 6 – stringency 3 --clip_R1 5 --clip_R2 10” using trim-galore (v 0.6.6), cutadapt (v3.4) and fastqc (0.11.9) ^51,52^(https://github.com/s-andrews/FastQC). Comprehensive quality control reports were generated using multiqc (v1.28)^53^.

The bisulfite reference genome for the human genome hg38 (GRCh38.primary) was prepared using the command “bismark_genome_preparation” in Bismark (v0.22.3)^54^. The trimmed reads were then mapped to the bisulfite genome using the parameters “--bowtie2 --parallel 16 --bam ‘$HG38_BISMARK’ −1 ‘$[FASTQ_R1_TRIMMED]’ −2 ‘$[FASTQ_R2_TRIMMED]’” (bowtie2 v2.4.4; samtools v1.12)^55,56^. The bam file was processed with deduplicate_bismark with default parameters to remove duplicate reads. The deduplicated bam file was used to calculate methylation level at each cytosine site (CpG and Non-CpG) using Bismark_methylation_extractor with the parameters “-p --gzip --comprehensive --multicore 5 --merge_non_CpG --bedGraph --counts --buffer_size 200G --cytosine_report --genome_folder ‘$HG38_BISMARK’ --report”. The resulting genomewide Cytosine Reports were used as inputs for differential methylation analysis with methylKit^57^. To identify groups of genes exhibiting coordinated changes in methylation and who share baseline methylation levels across contexts, for each gene the baseline was calculated as the average across all cytosines in that context (mCG, hmCG, mCH, hmCH) (cov > 5), and the number of significant cytosines (qvalue < 0.05, cov > 5) increasing (positive meth.diff) or decreasing (negative meth.diff) was counted for each context. Given the variability in gene size the signed magnitude of change was calcualted as shown below. Genes with incomplete data across any context were excluded. The remaining gene and feature matrix was scaled to unit variance and clustered using k-means, with the number of clusters selected using within-cluster sum of squares, silhouette width and stability across repeated subsampling.

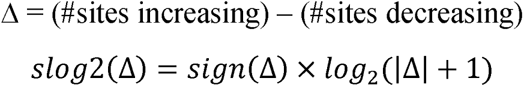

For visualization of methylation levels in IGV, .meth files were generated from the deduplicated bam files using methcounts and fasta file for HG38 with default parameters (Methpipe package (v5.0.1)^58,59^. These .meth files were used as inputs for the mlml tool (within the methpipe package) with the parameters “-v -u ‘$BS’ -m ‘$OXBS’” to estimate methylation and hydroxymethylation levels at each Cytosine position across the genome (separately for CpG and Non-CpG contexts)^60^. A custom R script was developed that dynamically split the MLML outputs, specifically, the mC and hmC data for each Cytosine and saved two separate bedGraph files. These bedGraphs were sorted by chromosome positions using bedSort (ucsc-bedsort-469) which were finally converted to bigwigs using bedgraphtobigwig (ucsc-bedgraphtobigwig-472) and visualized with IGV.

#### ATACseq Consensus peak calling and differential analysis

##### Consensus peak generation and filtering

NarrowPeak files from each sample were imported as *GRanges* objects via *rtracklayer::import()* (*rtracklayer).* To focus on summitcentered regions, each peak was resized to a 501 bp window (±250 bp) around its midpoint using *GenomicRanges::resize()* (*GenomicRanges*). Within each sample, we then applied an iterative clustering algorithm:

1. All peaks were merged into preliminary clusters via GenomicRanges::reduce().
2. Cluster membership was assigned by *findOverlaps()*, and within each cluster the single peak with the highest CPMnormalized score (computed from the original narrowPeak pvalue field using *edgeR::cpm())* was retained.
3. Retained peaks were removed from the pool, and steps 1–2 were repeated until no peaks remained.

The resulting samplespecific “winner” peaks were concatenated across all samples into a pansample GRangesList, unlisted, and sorted. We then built a peak × sample score matrix by reoverlapping each peak to every sample’s peak set (via findOverlaps()) and extracting CPM scores. Finally, peaks with CPM > 5 in at least two samples (i.e. rowSums(scoreMat > 5) ≥ 2) were selected to produce the final iterative consensus peak file filtered for 2 samples and at least 5CPM.

##### Read counting and differential analysis

The filtered consensus peaks were converted to a *SummarizedExperiment* object by counting overlaps between the iterative consensus peaks against all BAMs using *GenomicAlignments::summarizeOverlaps*. A DESeq2 dataset was constructed with *DESeqDataSetFromMatrix()* and tested for differential accessibility between experimental groups *(DESeq2::DESeq(); DESeq2)*, incorporating sample metadata from the design matrix. Variancestabilized counts were obtained via vst(), and principalcomponent plots were generated with plotPCA(). Differential peaks were called as padj < 0.05.

##### Peak Binarization and Cell type Assisgnment

To identify highly reproducible peaks selectively active in each cell type, we began with our final ATACseq SummarizedExperiment and sample metadata grouping samples by the Celltype column. Promoter regions were defined as ±150 bp around transcription start sites using GenomicRanges::promoters() on TxDb.Hsapiens.UCSC.hg38.knownGene Peaks overlapping these promoters were separated from nonpromoter peaks via findOverlaps(). Nonpromoter peaks were converted into a DESeq2 dataset and variance-stabilized counts computed via DESeq2::vst() yielding a matrix (vst_nonpromoter) in which rows are peaks and columns are samples. To facilitate comparisons across cell types, we reindexed rows sequentially and ensured row names matched the vst output. For each cell type with at least two replicates, we calculated the perpeak mean and standard deviation of vst counts across samples of that cell type using matrixStats::rowMeans2() and matrixStats::rowSds() The results were stored in matrices meansATAC and sdsATAC, with columns corresponding to cell types in a consistent order. To call binary presence/absence across cell types, we first sorted each peak’s means in ascending order (maintaining the matching cell type labels). For each cell type *i* (in rank order after the first), a peak was called “present” in *i* if

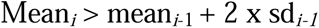

This threshold guards against calling peaks present in a cell type unless their signal exceeds the previous cell type’s mean by two standard deviations. Peaks called “present” for cell type *i* were removed from further consideration in subsequent rounds, and their binary membership vectors (indicating which of the remaining cell types they would also “pass” had they not been removed) were recorded. Combining these membership vectors across all iterations produces a perpeak, percelltype logical matrix. A concatenated label string (e.g. “Astrocyte_Oligo”) summarizes each peak’s cell type-specificity in a new clusterfull column. We merged this binary membership information back into the rowData of the nonpromoter DESeq2 object, aligning by peak index. The enriched rowData now includes, for each peak, logical flags for presence in each cell type and the composite clusterfull label.

##### software

– rtracklayer v1.58.0; GenomicRanges v1.50.1; GenomicAlignments v1.32.0; edgeR v3.38.4; SummarizedExperiment v1.30.0; DESeq2 v1.38.1; TxDb.Hsapiens.UCSC.hg38.knownGene v3.17.0; writexl v1.5.2; plus tidywrangling via stringr v1.5.0 and dplyr v1.1.2, matrixStats v0.62.0

#### cCRE mapping

In order to identify candidate cis-regulatory elements (cCREs) whose chromatin accessibility is linked to local gene expression, we developed a custom R function utilizing *rowCorr*, that integrates paired RNAseq and ATACseq data originating from the same tissue donor and brain region. Briefly, both assays are represented as *SummarizedExperiment* objects (SE_RNA and SE_ATAC), with rows corresponding to genes and peaks, respectively, and columns to paired biological samples. For each gene in SE_RNA, we define a genomic search window of ±1 Mb (configurable via the max.dist argument) around its transcription start site by resizing the gene ranges, and we pair these to ATAC seq peaks whose midpoints fall within that window. The resulting overlaps are recorded in a data frame *o* that includes the gene index, peak index and linear distance between each gene–peak pair.

Within *getCorr*, we then compute correlations of normalized accessibility and expression values for all gene–peak pairs using our C++ backed *rowCorCpp* routine, yielding a vector of correlation coefficients per pair. Test statistics are converted from correlation to t statistics accounting for the number of samples, and two sided P values are derived via the Student’s t distribution. False discovery rates (FDR) are controlled by the Benjamini–Hochberg procedure to produce FDR adjusted P values.

To assess background correlation levels, the auxiliary function .*getNullCorrelations* constructs a chromosome stratified null distribution: for each chromosome, peaks on all other chromosomes are randomly sampled (up to 1,000 per chromosome) and correlated against genes on the focal chromosome to generate a pool of “trans” correlations. The empirical mean and standard deviation of this null pool are then used to compute empirical P values for the observed cis correlations, with subsequent FDR correction. Finally, gene–peak pairs with both nominal and empirical FDR below predefined thresholds can be labeled as cCREs for downstream analysis.

#### FANS-C&R-seq Nuclear Staining, Library Preparation and Processing

Nuclei were isolated from fresh frozen brain tissue samples as previously described in ***Basic Protocol 1***^21^, but with modified fixation conditions to ensure sensitive detection of C&R-seq targets. Once isolated, bulk nuclei were lightly fixed with 0.5% methanol-free formaldehyde for 1 minute rotating at RT, quenched with 125mM glycine for 5 minutes rotating, washed, then resuspended in 1ml FANS wash buffer (1X PBS, 0.5% BSA, 1X Roche cOmplete EDTA-free Protease Inhibitor) with 0.01% digitonin to block and permeabilize for 30 minutes rotating at RT. 10µl of nuclei were removed for yield quantification via Thermo Fisher Countess II, then sample was partitioned into 5 pools of nuclei for 5 separate histone PTM target conditions – euchromatic marks H3K4me3 (Active Motif 39159) and H3K27ac (Active Motif 39133), and rabbit IgG negative control (Epicypher 13-0042). FANS-C&R antibody buffer (FANS wash buffer, 0.01% digitonin, 0.5mM Spermidine, 2mM EDTA) was then prepared containing appropriate FANS labelling antibody dilutions as described in ***Basic Protocol 3*** (striatal MSNs) ^21^and in ^5^ (cerebellar granule neurons and glial cell types of the striatum).

Partitioned bulk nuclei conditions were pelleted (4 minutes, 800xg). Due to yield variability between donor aliquots, the following guidelines for bulk nuclei resuspension were established to obtain reproducible target signals, each target condition must contain: 1) 3-4x10^3 nuclei/µl of FANS-C&R-antibody buffer, 2) a minimum of 50µl FANS-C&R-antibody buffer per target condition, 3) 0.01 ug/µl histone PTM target antibody. Once resuspended, each condition was incubated rotating 4C overnight. Samples were then processed for secondary antibody labeling and sorted into neuronal and glial cell types for each histone PTM target condition as described in ***Basic Protocol 3*** ^21^ and in^61^. Sorted nuclei counts for each histone PTM condition and cell type sample were kept within the range of 1x10^4 – 3x10^5, with maximum yield prioritized due to sample scarcity.

C&R-seq was carried out same-day using a modified protocol from (Meers et al. 2019) and (Epicypher C&R Manual v5.1) with reagents from CUTANA™ ChIC / CUT&RUN kit (Epicypher 14-1048). Sorted samples were centrifuged (4 minutes, 800xg), resuspended in C&R-wash buffer (20mM HEPES pH 7.5, 150mM NaCl, 0.5mM Spermidine, 1x Roche cOmplete EDTA-free Protease Inhibitor), centrifuged again, resuspended in 100µl C&R-wash buffer and transferred to labeled 8-strip tubes. 10.5µl of activated ConA beads (see Epicypher C&R Manual 5.1 for activation protocol) were added to each 100µl reaction and incubated for 15 minutes RT on a gentle lateral nutator. Sample 8-strip tubes were placed on a 96 tube magnetic rack (Thermo Fisher AM10050) for 3 minutes, supernatant was removed and replaced with 200µl cold C&R-wash buffer while still on magnet. Samples were washed twice more in cold C&R-DIG buffer (C&R-wash buffer, 0.01% digitonin), then resuspended in 50µl cold C&R-DIG buffer with 1X pA/G-MNase (Epicypher, 15-1016) and incubated 10 minutes RT on a gentle lateral nutator. After returning to magnetic rack on ice and washing three times in cold C&R-DIG buffer, samples were resuspended in 50µl cold C&R-DIG buffer containing 2mM Cacl2 and incubated rotating at 45 ° angle for 2 hours at 4C to cleave pA/G-Mnase-bound chromatin targets. 34µl of RT C&R-Stop buffer (340mM NaCl, 20mM EDTA, 4 mM EGTA, 50µg/mL RNase A, 50ug/mL Glycogen) containing 1pg E.Coli K12 spike-in DNA (Epicypher 18-1401) was added to samples, which were then incubated in thermocycler for 10 min at 37C to release cleaved chromatin into supernatant. Nuclei were quick-centrifuged and sample chromatin supernatants were removed after a 3 minute incubation on magnet. 1µl 8% SDS and 2µl 10µg/µl Proteinase K (Qiagen, 19131) were added to all extracted sample chromatin supernatants, which were then mixed and incubated overnight at 55C via thermocycler.

The following day, samples were transferred to low-bind microcentrifuge tubes, brought up to 250µl with water, then vortexed with 250µl phenol:chloroform:isoamyl alcohol 25:24:1 pH 8 (Sigma-Aldrich P2069) and incubated 10 mins RT. Samples were quick-centrifuged, then transferred to pre-centrifuged phase lock gel heavy tubes (Qiagen, 129056) and centrifuged (5 min, 16000xg). 250µl chloroform (Sigma-Aldrich C2432) was mixed with sample aqueous phases, and tubes were centrifuged again. Sample aqueous phases were transferred to new pre-centrifuged phase lock gel heavy tubes, mixed with 400µl chloroform and centrifuged (3 minutes, 12000xg). Sample aqueous phases were transferred to new microcentrifuge tubes and brought up to 250µl with water. Samples were vortexed with 25µl 3M NaAc (Thermo Fisher R1181), 550µl 100% EtOH, and 15µg GlycoBlue™ Coprecipitant (Thermo Fisher AM9515) and placed at −20C for 1-4 days to precipitate cleaved DNA. Samples were then centrifuged down into DNA pellets (1 hour, 20000xg 4C), washed twice in 1ml 75% EtOH (5 minutes, 20000xg), air dried for 5-10 minutes, then DNA was resuspended in 25µl 0.1X TE buffer and frozen.

Libraries were prepared from purified sample DNA using the CUTANA™ CUT&RUN Library Prep Kit (Epicypher 14-1001, 14-1002) as described by (Epicypher C&R library prep manual v1.5) with the following modifications: 1) Illumina adapter was reduced to 0.75 pmol per sample, 2) index amplification was raised to 16 cycles, 3) sample libraries were brought up to 50µl with 0.1X TE before the second bead cleanup, 4) the final elution volume was raised to 14µl 0.1X TE with 13µl sample library extracted. Due to an increased likelihood of adapter-dimer formation caused by low sample input material, fragment size distribution was quantified on the Agilent 4200 TapeStation system using 1µl of each sample library. The molar proportion of dimer fragments (0-170bp) to signal fragments (170-1000bp) was used to pool all samples on a weighted basis, producing a pool containing equimolar signal fragments for each sample. Pool underwent 1X SPRI bead clean up (Beckman Coulter) and was sequenced 50-80million reads per sample on NovaSeq6000 (2 × 100 bp).

Sample paired-end FastQs were aligned in R using rsubread::align() with fragment length range of 10bp-600bp, first aligned against human genome GRCh38.p13 and then aligned against E.Coli MG1655 for downstream spike-in normalization. Peaks were called in python using MACS2 callpeak function with flags -f BAMPE -p 1e-5 --keep-dup=all -g hs -c <appropriate IgG control>.

*Software: Rsubread 2.16.0, macs2 2.2.9.1*

#### In situ hybridization (ISH) and image analysis

ISH was performed using RNAscope^TM^ Multiplex Fluorescent Reagent Kit v2 (Advanced Cell Diagnostics, 323100) according to the manufacturer’s instructions for fresh-frozen samples with the following modifications: sections were fixed for 15 minutes in chilled 4% formaldehyde in PBS, then rinsed in PBS, dehydrated, and photobleached in 100% ethanol for 40-45 hours using a photobleaching device^62^. The following probes were used: Hs COCH (Advanced Cell Diagnostics, 1104401-C1), Hs PDE10A (Advanced Cell Diagnostics, 466151-C2), Hs PCP4 (Advanced Cell Diagnostics, 446111-C2), TSA Vivid^TM^ fluorophore 570 (Advanced Cell Diagnostics, #323272, used for COCH detection) and TSA Vivid^TM^ fluorophore 650 (Advanced Cell Diagnostics, #323273, used to detect PDE10A and PCP4 probes). Sections were counterstained with DAPI and then cover-slipped with ProLong™ Diamond Antifade Mountant (ThermoFisher, P36970). Images were acquired on a Zeiss Confocal Microscope LSM 710 confocal microscope using a Plan-Apochromat 20x, 0.8 NA objective lens.

Regions of interest (ROIs), each containing an MSN, were defined in ImageJ (1.54f) based solely on the signals from DAPI and the probe for *COCH*, an MSN marker gene whose expression level does not change in HD based on our previously published data^5^. For each ROI, the number of *COCH*+ puncta and *PDE10A*+ puncta (84-305 ROIs per donor) or *PCP*+ puncta (38-224 ROIs per donor) was counted from each ROI using a previously published ImageJ script (https://zenodo.org/records/14506480)^63^. The ‘*PDE10A* puncta’/’*COCH* puncta’ or ‘*PCP4* puncta’/’*COCH* puncta’ ratios were calculated and plotted separately for each donor. Each donor’s median value of these ratios was used for statistical analysis of the difference between HD and control donors.

## Notes

### Competing Interest Statement

The authors have declared no competing interest.

### Summary of Updates

Figures 2-5 were redone and the manuscript was rewritten for clarity.

## References

1. A novel gene containing a trinucleotide repeat that is expanded and unstable on Huntington’s disease chromosomes. The Huntington’s Disease Collaborative Research Group. Cell 72, 971–83 (1993).

2. Kennedy, L. et al. Dramatic tissue-specific mutation length increases are an early molecular event in Huntington disease pathogenesis. Hum Mol Genet 12, 3359–67 (2003).

3. Handsaker, R.E. et al. Long somatic DNA-repeat expansion drives neurodegeneration in Huntington’s disease. Cell 188, 623–639 e19 (2025).

4. Pressl, C. et al. Selective vulnerability of layer 5a corticostriatal neurons in Huntington’s disease. Neuron 112, 924–941 e10 (2024).

5. Matlik, K. et al. Cell-type-specific CAG repeat expansions and toxicity of mutant Huntingtin in human striatum and cerebellum. Nat Genet 56, 383–394 (2024).

6. Genetic Modifiers of Huntington’s Disease, C. Genetic modifiers of somatic expansion and clinical phenotypes in Huntington’s disease highlight shared and tissue-specific effects. Nat Genet 57, 1426–1436 (2025).

7. Genetic Modifiers of Huntington’s Disease Consortium. Electronic address, g.h.m.h.e. & Genetic Modifiers of Huntington’s Disease, C. CAG Repeat Not Polyglutamine Length Determines Timing of Huntington’s Disease Onset. Cell 178, 887–900 e14 (2019).

8. Genetic Modifiers of Huntington’s Disease, C. Identification of Genetic Factors that Modify Clinical Onset of Huntington’s Disease. Cell 162, 516–26 (2015).

9. Wang, N. et al. Distinct mismatch-repair complex genes set neuronal CAG-repeat expansion rate to drive selective pathogenesis in HD mice. Cell 188, 1524–1544 e22 (2025).

10. Mouro Pinto, R., et al. In vivo CRISPR-Cas9 genome editing in mice identifies genetic modifiers of somatic CAG repeat instability in Huntington’s disease. Nat Genet 57, 314–322 (2025).

11. Wheeler, V.C. & Dion, V. Modifiers of CAG/CTG Repeat Instability: Insights from Mammalian Models. J Huntingtons Dis 10, 123–148 (2021).

12. Kovalenko, M. et al. Msh2 acts in medium-spiny striatal neurons as an enhancer of CAG instability and mutant huntingtin phenotypes in Huntington’s disease knock-in mice. PLoS One 7, e44273 (2012).

13. Gu, X. et al. Uninterrupted CAG repeat drives striatum-selective transcriptionopathy and nuclear pathogenesis in human Huntingtin BAC mice. Neuron 110, 1173–1192 e7 (2022).

14. Horvath, S. et al. Huntington’s disease accelerates epigenetic aging of human brain and disrupts DNA methylation levels. Aging (Albany NY) 8, 1485–512 (2016).

15. Lee, H. et al. Cell Type-Specific Transcriptomics Reveals that Mutant Huntingtin Leads to Mitochondrial RNA Release and Neuronal Innate Immune Activation. Neuron 107, 891–908 e8 (2020).

16. Langfelder, P. et al. Integrated genomics and proteomics define huntingtin CAG length-dependent networks in mice. Nat Neurosci 19, 623–33 (2016).

17. Merienne, N. et al. Cell-Type-Specific Gene Expression Profiling in Adult Mouse Brain Reveals Normal and Disease-State Signatures. Cell Rep 26, 2477–2493 e9 (2019).

18. Achour, M. et al. Neuronal identity genes regulated by super-enhancers are preferentially down-regulated in the striatum of Huntington’s disease mice. Hum Mol Genet 24, 3481–96 (2015).

19. Brule, B. et al. Accelerated epigenetic aging in Huntington’s disease involves polycomb repressive complex 1. Nat Commun 16, 1550 (2025).

20. Alcala-Vida, R. et al. Age-related and disease locus-specific mechanisms contribute to early remodelling of chromatin structure in Huntington’s disease mice. Nat Commun 12, 364 (2021).

21. Pressl, C., et al. Isolation and Molecular Profiling of Nuclei of Specific Neuronal Types from Human Cerebral Cortex and Striatum. Curr Protoc 4, e70067 (2024).

22. Li, Y.E. et al. An atlas of gene regulatory elements in adult mouse cerebrum. Nature 598, 129–136 (2021).

23. Zhang, K. et al. A single-cell atlas of chromatin accessibility in the human genome. Cell 184, 5985–6001 e19 (2021).

24. Granja, J.M. et al. ArchR is a scalable software package for integrative single-cell chromatin accessibility analysis. Nat Genet 53, 403–411 (2021).

25. Loupe, J.M. et al. Promotion of somatic CAG repeat expansion by Fan1 knock-out in Huntington’s disease knock-in mice is blocked by Mlh1 knock-out. Hum Mol Genet 29, 3044–3053 (2020).

26. Bunting, E.L. et al. Antisense oligonucleotide-mediated MSH3 suppression reduces somatic CAG repeat expansion in Huntington’s disease iPSC-derived striatal neurons. Sci Transl Med 17, eadn4600 (2025).

27. Creyghton, M.P. et al. Histone H3K27ac separates active from poised enhancers and predicts developmental state. Proc Natl Acad Sci U S A 107, 21931–6 (2010).

28. Wang, D. et al. Active DNA demethylation promotes cell fate specification and the DNA damage response. Science 378, 983–989 (2022).

29. Adachi, K. et al. Esrrb Unlocks Silenced Enhancers for Reprogramming to Naive Pluripotency. Cell Stem Cell 23, 900–904 (2018).

30. Stoyanova, E., Riad, M., Rao, A. & Heintz, N. 5-Hydroxymethylcytosine-mediated active demethylation is required for mammalian neuronal differentiation and function. Elife 10(2021).

31. He, Y.F. et al. Tet-mediated formation of 5-carboxylcytosine and its excision by TDG in mammalian DNA. Science 333, 1303–7 (2011).

32. Tahiliani, M. et al. Conversion of 5-methylcytosine to 5-hydroxymethylcytosine in mammalian DNA by MLL partner TET1. Science 324, 930–5 (2009).

33. Mellen, M., Ayata, P. & Heintz, N. 5-hydroxymethylcytosine accumulation in postmitotic neurons results in functional demethylation of expressed genes. Proc Natl Acad Sci U S A 114, E7812–E7821 (2017).

34. Whyte, W.A. et al. Master transcription factors and mediator establish super-enhancers at key cell identity genes. Cell 153, 307–19 (2013).

35. Appenzeller, S. et al. Autosomal-dominant striatal degeneration is caused by a mutation in the phosphodiesterase 8B gene. Am J Hum Genet 86, 83–7 (2010).

36. Butler, K.M. et al. De novo variants in GABRA2 and GABRA5 alter receptor function and contribute to early-onset epilepsy. Brain 141, 2392–2405 (2018).

37. Diggle, C.P. et al. Biallelic Mutations in PDE10A Lead to Loss of Striatal PDE10A and a Hyperkinetic Movement Disorder with Onset in Infancy. Am J Hum Genet 98, 735–43 (2016).

38. Justice, J.L. et al. Multi-epitope immunocapture of huntingtin reveals striatum-selective molecular signatures. Mol Syst Biol 21, 492–522 (2025).

39. Goldstrohm, A.C., Albrecht, T.R., Sune, C., Bedford, M.T. & Garcia-Blanco, M.A. The transcription elongation factor CA150 interacts with RNA polymerase II and the pre-mRNA splicing factor SF1. Mol Cell Biol 21, 7617–28 (2001).

40. Dragileva, E. et al. Intergenerational and striatal CAG repeat instability in Huntington’s disease knock-in mice involve different DNA repair genes. Neurobiol Dis 33, 37–47 (2009).

41. Holbert, S. et al. The Gln-Ala repeat transcriptional activator CA150 interacts with huntingtin: neuropathologic and genetic evidence for a role in Huntington’s disease pathogenesis. Proc Natl Acad Sci U S A 98, 1811–6 (2001).

42. Guo, J.U. et al. Distribution, recognition and regulation of non-CpG methylation in the adult mammalian brain. Nat Neurosci 17, 215–22 (2014).

43. Sonn, J.Y. et al. MeCP2 interacts with the super elongation complex to regulate transcription. Sci Adv 11, eadt5937 (2025).

44. Azuma, R. et al. A novel mutation of PDE8B Gene in a Japanese family with autosomal-dominant striatal degeneration. Mov Disord 30, 1964–7 (2015).

45. Srivastava, S. et al. Loss-of-function variants in HIVEP2 are a cause of intellectual disability. Eur J Hum Genet 24, 556–61 (2016).

46. Rahimi, M.J. et al. De novo variants in ATP2B1 lead to neurodevelopmental delay. Am J Hum Genet (2025).

47. Charlesworth, G. et al. Mutations in ANO3 cause dominant craniocervical dystonia: ion channel implicated in pathogenesis. Am J Hum Genet 91, 1041–50 (2012).

48. Winkelmann, J. et al. Mutations in DNMT1 cause autosomal dominant cerebellar ataxia, deafness and narcolepsy. Hum Mol Genet 21, 2205–10 (2012).

49. Love, M.I., Hogenesch, J.B. & Irizarry, R.A. Modeling of RNA-seq fragment sequence bias reduces systematic errors in transcript abundance estimation. Nat Biotechnol 34, 1287–1291 (2016).

50. Love, M.I., Soneson, C. & Patro, R. Swimming downstream: statistical analysis of differential transcript usage following Salmon quantification. F1000Res 7, 952 (2018).

51. Martin, M. Cutadapt removes adapter sequences from high-throughput sequencing reads. EMBnet. journal 17, 10–12 (2011).

52. Krueger, F. Trim galore. A wrapper tool around Cutadapt and FastQC to consistently apply quality and adapter trimming to FastQ files 516, 517 (2015).

53. Ewels, P., Magnusson, M., Lundin, S. & Kaller, M. MultiQC: summarize analysis results for multiple tools and samples in a single report. Bioinformatics 32, 3047–8 (2016).

54. Krueger, F. & Andrews, S.R. Bismark: a flexible aligner and methylation caller for Bisulfite-Seq applications. Bioinformatics 27, 1571–2 (2011).

55. Li, H. et al. The Sequence Alignment/Map format and SAMtools. Bioinformatics 25, 2078–9 (2009).

56. Langmead, B. & Salzberg, S.L. Fast gapped-read alignment with Bowtie 2. Nat Methods 9, 357–9 (2012).

57. Akalin, A. et al. methylKit: a comprehensive R package for the analysis of genome-wide DNA methylation profiles. Genome Biol 13, R87 (2012).

58. Robinson, J.T. et al. Integrative genomics viewer. Nat Biotechnol 29, 24–6 (2011).

59. Song, Q. et al. A reference methylome database and analysis pipeline to facilitate integrative and comparative epigenomics. PLoS One 8, e81148 (2013).

60. Qu, J., Zhou, M., Song, Q., Hong, E.E. & Smith, A.D. MLML: consistent simultaneous estimates of DNA methylation and hydroxymethylation. Bioinformatics 29, 2645–6 (2013).

61. Matlik, K., Abo-Ramadan, U., Harvey, B.K., Arumae, U. & Airavaara, M. AAV-mediated targeting of gene expression to the peri-infarct region in rat cortical stroke model. J Neurosci Methods 236, 107–13 (2014).

62. Murakami, T.C. et al. Open-source Photobleacher for Fluorescent Imaging of Large Pigment-Rich Tissues. bioRxiv, 2025.02.24.639965 (2025).

63. Zou, Y. et al. ATG8 delipidation is not universally critical for autophagy in plants. Nat Commun 16, 403 (2025).

